# Systematic analysis and optimization of early warning signals for critical transitions

**DOI:** 10.1101/2022.11.04.515178

**Authors:** Daniele Proverbio, Alexander Skupin, Jorge Gonçalves

**Affiliations:** Luxembourg Centre for Systems Biomedicine; University of Luxembourg; Belvaux, 6 Avenue du Swing, 4367; Luxembourg; College of Engineering, Mathematics and Physical Sciences; University of Exeter; Exeter, EX4 4QL; UK; National Center for Microscopy and Imaging Research; University of California San Diego; La Jolla, Gilman Drive, CA, 9500; United States; Department of Physics and Material Science; University of Luxembourg; Luxembourg, 162a Avenue de la Faiencerie, 1511; Luxembourg; Department of Plant Sciences; University of Cambridge; Cambridge, CB2 3EA; UK

**Keywords:** Critical transition, Early warning signals, Bistable systems, Optimisation, Multivariate analysis, Bifurcation, Dynamics

## Abstract

Abrupt shifts between alternative regimes occur in complex systems, from cell regulation to brain functions to ecosystems. Several model-free Early Warning Signals (EWS) have been proposed to detect impending transitions, but failure or poor performance in some systems have called for better investigation of their generic applicability. In particular, there are still ongoing debates whether such signals can be successfully extracted from data. In this work, we systematically investigate properties and performance of dynamical EWS in different deteriorating conditions, and we propose an optimised combination to trigger warnings as early as possible, eventually verified on experimental data. Our results explain discrepancies observed in the literature between warning signs extracted from simulated models and from real data, provide guidance for EWS selection based on desired systems and suggest an optimised composite indicator to alert for impending critical transitions.

**Highlights:**

- How to extract early warning signals (EWS) against critical transitions from data is still poorly understood
- A mathematical framework assesses and explains the performance of EWS in noisy deteriorating conditions
- Composite indicators are optimised to alert for impending shifts
- The results are applicable to wide classes of systems, as shown with models and on empirical data.

## 1. Introduction

The dynamics of many complex systems is characterised by critical thresholds (tipping points) and abrupt shifts between alternative regimes (Scheffer et al., 2009; Ashwin and Zaikin, 2015). Various examples have been observed in diverse research fields and include collapses of ecosystems (Hirota et al., 2011; Wang et al., 2012a), sudden climate shifts (Lenton et al., 2012; Drijfhout et al., 2015) or financial crashes (Dmitriev et al., 2017; Diks et al., 2019). Abrupt regime shifts have particularly been theorised and observed in systems biology and medicine (Korolev et al., 2014; Trefois et al., 2015; Aihara et al., 2022), at the onset of certain disease states like atrial fibrillation (Quail et al., 2015) or epileptic seizures (Meisel and Kuehn, 2012), as well as in biological processes like regulation of gene networks (Angeli et al., 2004; Sharma et al., 2016) and cell fate decisions (Ghaffarizadeh et al., 2014; Mojtahedi et al., 2016), including epithelial-mesenchymal transitions (Lang et al., 2021). Correctly detecting and alerting for these critical changes allows to better understand complex developments and to anticipate dangerous outcomes. However, many such complex systems have not been fully characterised with mechanistic models, thus requiring simpler and more generic approaches to support data-driven estimates.

The critical transitions (CT) framework have been proposed to address tipping points using low-dimensional systems descriptions (Kuehn, 2011) and associated early warning signals (EWS), computed from statistical indicators extracted from data like increasing variance, autocorrelation or coefficient of variation (Drake and Griffen, 2010; Lade and Gross, 2012). These signs and derived indexes (Chen et al., 2012; Navid Moghadam et al., 2020; Matsumori et al., 2019), in principle generic for broad classes of systems, have been tested and applied on biological, epidemiological and medical data with alternate success (Carpenter et al., 2011; Dai et al., 2012; Wilkat et al., 2019; Proverbio et al., 2022a). Therefore, recent studies have recommended caution when attempting predictions based on EWS (Boettiger and Hastings, 2012; Clements and Ozgul, 2018; Dudney and Suding, 2020). Since there is an increasing interest for EWS in systems biology and biomedicine, it is thus compelling to provide a unified framework for the analysis and interpretation of such indicators, to determine in which cases they can be safely applied and to understand their limitations. In addition, going beyond univariate indicators will improve their performance in detecting and alerting for impending critical transitions.

In this work, we provide a systematic analysis of the CT framework and its associated EWS, to define their range of applicability and understand why discrepancies have been observed between theoretical predictions and experimental data (Kuehn et al., 2022; Cohen et al., 2022). Systems biology is characterised by two main paradigms (Mazzocchi, 2012): one investigating the single details of molecular combinations or regulatory networks, alike to “microstates” in statistical mechanics (Stumpf et al., 2017), and another looking for general analytical models, built upon kinetic theories, to understand complicated biochemical processes in simpler and general terms (Ferrell Jr et al., 2009). The latter allows to construct classes of systems according to universal routes of dynamical development, regardless of the microscopic details. We leverage this paradigm to make sense of critical transitions and identify the most relevant classes pertaining to biological systems (Box 1). We also provide guidance for EWS selection and optimisation, depending on realistic noise properties and other notable features of classes of complex systems, developing new composite indicators.

Our work bridges mathematical insights and observations of real systems to classify various tipping mechanisms. There are ongoing debates whether regime shifts in biological systems, like cell-fate decision, are primarily driven by deterministic bifurcations (Andrecut et al., 2011; Stanoev et al., 2021) or by random fluctuations (Wang et al., 2011; Stumpf et al., 2017), which prompted several authors to question the old “Waddington landscape” interpretation (Moris et al., 2016). By systematically analysing known regime shifts, we classify the mathematical models to address various types of critical transitions, subject to combinations of bifurcations and noise (Berglund and Gentz, 2006), and to develop a method to extract systems’ robustness proxies from data (Box 2).

We first employ a framework based on dynamical manifolds, underpinning universal routes to explosive transitions (Kuehn and Bick, 2021), to characterise the warning signals associated to “noisy” bifurcations, and to study their dependency on noise properties and other dynamical features like rapid approaches to threshold values. This way, we provide general results about EWS robustness and sensitivity to dynamical features, to guide applications on various systems, understand their limitations and promote future developments. Then, we focus on a critical transition sub-class of high biological relevance, the stochastic saddle-node bifurcation (Ferrell Jr et al., 2009). For this tractable, yet realistic model of complex biological processes, we develop a composite EWS indicator to optimise the leading time of the alerts, i.e., how much in advance reliable signals are triggered, with respect to an impending transition. The new indicator is optimised over realistic noise types using the common genetic toggle switch model (Sharma et al., 2016), as representative of the considered CT class. This way, we overcome the limitations of other EWS from literature, which have mostly been developed over Gaussian noise while biological systems usually feature correlated and state-dependent noise (Hasty et al., 2000; Dunlop et al., 2008; Zhang et al., 2012). Thanks to this extension, the indicator also provides additional insights about the systems under investigation, such as inference of noise type from data. The theoretical results are finally tested and verified on publicly available experimental data, demonstrating their potential for monitoring and interpreting diverse systems.

### Box 1

**Classification of critical transitions**

Consider a dynamical system whose state (or regime) is usefully characterised by a set of dynamic variables *x* ∈ ℝ^*n*^, whose relations to each other are modeled by a set of parameters *p* ∈ ℝ^*m*^

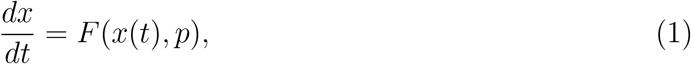

where *F* : ℝ^*n*+*m*^ → ℝ^*n*^ is a system of sufficiently smooth functions. If *p* is not explicitly dependent on time, the system is termed *autonomous*; if *p* = *p*(*t*), the system is called *non-autonomous*. The distinction between autonomous and non-autonomous can be supported when considering naturally fixed parameters (Maini et al., 1991), or when addressing timescale separation (“slow-fast system”) between biochemical processes, like mRNA transcription versus protein degradation times (Yasemi and Jolicoeur, 2021). This results in sets of dynamical (for variables) and algebraic (for parameters, termed at quasi-steady state) equations (Del Vecchio et al., 2016). Together, variables and parameters define and shape a state space (or “landscape”) that, if *F* (*x, p*) has elements of non-linearity, can be characterised by multiple attractors (MacArthur et al., 2009), i.e., region of stability for systems’s states. If parameters are allowed to change (either non-autonomously, or at quasi-steady state), the state space is dynamic and attractors can change, as opposed to static landscapes like Waddington’s.

The state space can be multidimensional. However, near bifurcation points, it can be aptly described using low-dimensional models associated to critical thresholds in the values of leading parameters (usually corresponding to the largest eigenvalues (Kuznetsov, 2013)). Such models are termed “normal forms” of a dynamical system, simplified minimal-order forms that determine the system’s behaviour and retain universal properties of generic bifurcations (see Kuehn and Bick (2021) and STAR Method C.1). Normal forms can be inferred from bistability properties (Angeli et al., 2004) or deduced from network models, if they are available for the considered systems (Gao et al., 2016; Tu et al., 2021).

In addition to bifurcation points, noise can characterise the system’s dynamics. Noise is ubiquitous in biology (Tsimring, 2014; Su et al., 2019) and can correspond to stochasticity in intrinsic biochemical processes or cell-cell variation (Zhang et al., 2012). Mathematically, noise variables can be modelled as fast degrees of freedom augmenting system (1), which is a dualistic representation to stochastic processes (Berglund and Gentz, 2006). Noise can push the system out of original attractors onto new ones, therefore causing random switches between phenotypic states even in the absence of dynamical bifurcations.

We propose to use the relative timescales between dynamical variables, parameters and noise to develop a systematic classification of transitions between system states. This way, we synthesise and improve the contributions of Thompson and Sieber (2011); Kuehn (2011); Ashwin et al. (2012); Shi et al. (2016) towards the establishment of a theory on critical transitions in real systems. To do so, extend and disentangle Eq. 1 to explicit the dependencies on state variables *x*∈ ℝ^*m*^ and system parameters *p* ∈ ℝ^*n*^, on the introduced stochastic variables *ξ* ∈ ℝ^*l*^ and on the relative timescales modelled by time parameters *τ*_*i*_, *i* = {*x, p, ξ*}. This results in a multiscale slow-fast system

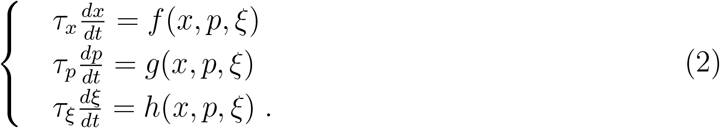

Using this representation, tipping systems can be classified into three main classes of critical transitions on the basis of relative timescales: bifurcation-induced (“b-tipping”), noise-induced (“n-tipping”) and rate-induced (“r-tipping”), following the nomenclature introduced by Ashwin et al. (2012):

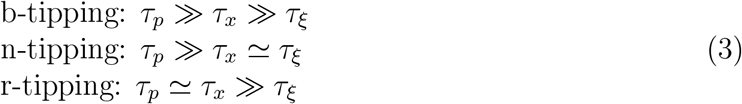

If *τ*_*ξ*_ *> τ*_*x*_, the system becomes ergodic and visits the full state-space uniformly without displaying transitions (Shi et al., 2016).

The b-tipping class thus encompasses all those transitions primarily driven by bifurcations, i.e., slow changes in control mechanisms modelled as quasi-steady approaches of leading parameters to their threshold values. They modify the attractor landscape, in the presence of low noise-to-signal ratios, and can be further sub-classified according to dimension *m* and co-dimension *n* (Thompson and Sieber, 2011). In this work, we only consider lowdimensional ones, commonly found in cell dynamics studies. Examples include toggle-switch mechanisms for the lac-operon (Ozbudak et al., 2004), population collapses of microbiological colonies past threshold concentrations of stressors or nutrients (Dai et al., 2015), or epithelial-mesenchymal determination (Sarkar et al., 2019). Higher *m* and *n* yield more complex bifurcations associated to, e.g., neural network activity (Izhikevich, 2007).

The n-tipping class groups various transitions driven by stochastic fluctuations on fixed landscapes, including large, impactful and unexpected events (sometimes called “dragon kings” (Sornette, 2006)). Example range from enzymes crossing activation chemical barriers via “promoting vibrations” (Antoniou and Schwartz, 2011), “rebellious cells” undergoing contrasting development pathways during cell reprogramming (Mojtahedi et al., 2016), and other long-studied cases of noise-induced transitions (Horsthemke and Lefever, 1984).

B-tipping and n-tipping directly link to the the aforementioned debates in systems biology about deterministic or stochastic drivers of critical changes. R-tipping refers to critical ramping of control parameters, not coped by the system, which has been so far observed in climate (Wieczorek et al., 2011) and engineering (Bonciolini et al., 2018) systems. The heat-shock response of plants to ramping temperature conditions (Moejes et al., 2017) may fall within this class, but further studies are required. The critical transition classes can be visualised on bifurcation diagrams or using quasi-potential landscapes (Zhou et al., 2012), which can be obtained as integrals of vector fields like Eq. 1 or inferred from data.

Fig. 1 shows the classification between the transition classes, with illustrative examples of what can happen to systems within simplified attractors. Note that the hard-cut classification derives from the mathematical assumptions in Eq. 3: gradients between the transition classes may exist and call for deep investigation. In particular, our work focuses on “noisy bifurcations”, i.e., dynamics characterised by bifurcation points and the presence of low to moderate noise-to-signal ratio.

**Figure 1:**
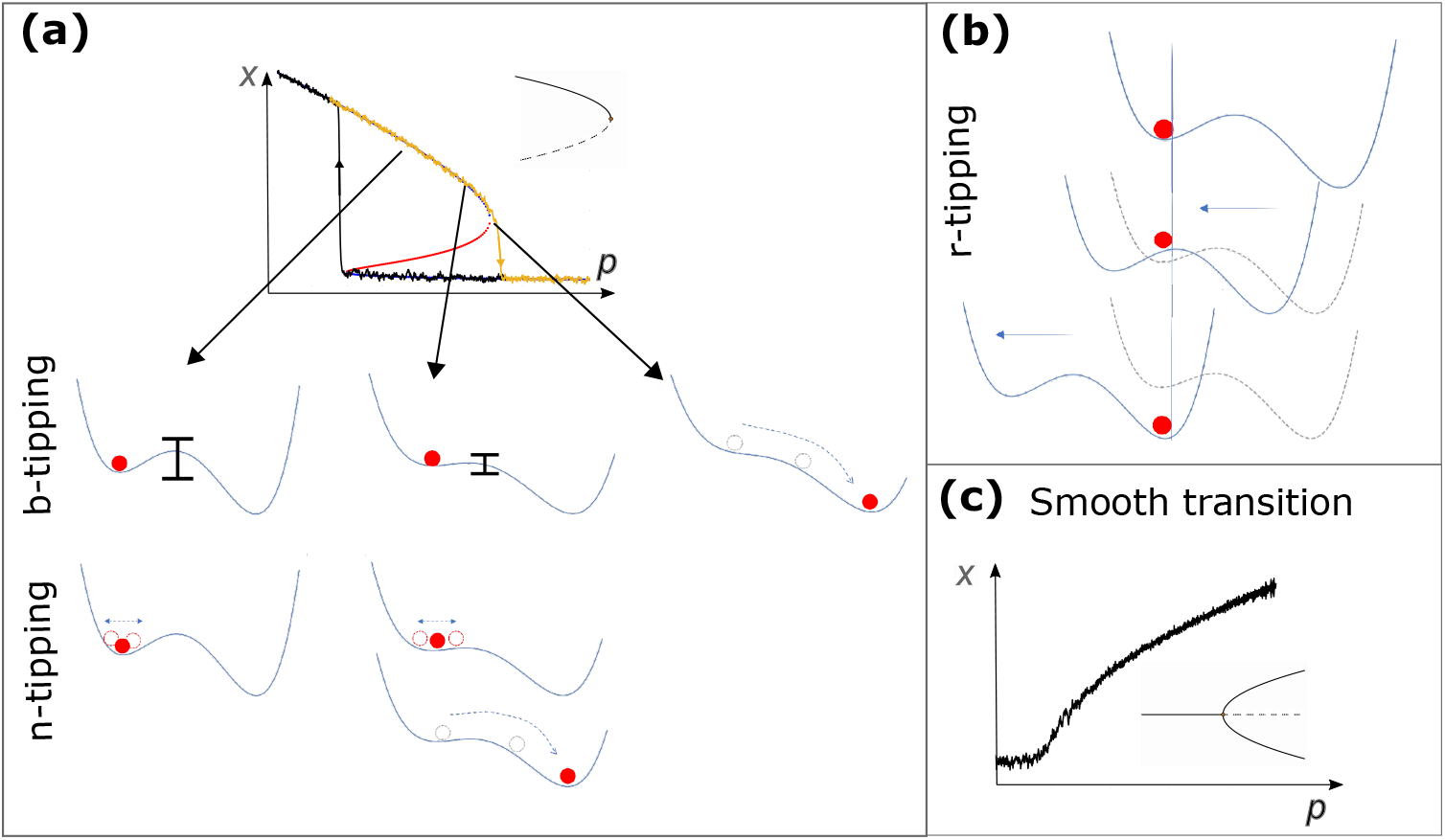
Classification of transitions between states *x* of a dynamical system, controlled by a slow changing parameter. *p. x* and *p* may also correspond to network combinations of variables and parameters (Moris et al., 2016; Gao et al., 2016). (a): Illustration of b-tipping and n-tipping using a bistable system with saddle-node bifurcations (unstable branch in red; saddle-node template shown in inset). Hysteresis can occur, i.e., asymmetric routes to tipping from one stable state or from the other (orange, from up to down with increasing *p*; black, from down to up with decreasing *p*). B-tipping: the system approaches the bifurcation point. The associated landscape is molded by *p* and the basin of attraction becomes shallower (as visualised by the bars) until disappearing; there, the system tips. N-tipping: if subject to strong fluctuations, depicted as wiggling of the red ball, the system can be pushed over the barrier onto an alternative attractor, even before the bifurcation point. (b) Illustration of r-tipping: rapid ramping of the control parameter makes it as if the landscape shifts and the systems does not manage to move along, therefore tipping onto another attractor “sliding” underneath. See Ashwin et al. (2012) for formal definitions. (c): Example of “smooth” transition without hysteresis, using a dynamical system close to a pitchfork bifurcation (inset) as template. To reproduce the plots, see STAR Method C.3.

### Box 2

**Bifurcations with noise and system robustness**

Among the critical transition classes described above, let us consider those primarily driven by bifurcations, with noise further influencing the dynamics. In this sense, we can speak of “noisy b-tipping”, with the first condition in Eq. 3 becoming

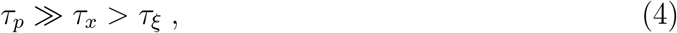

that is, the noise-to-signal ratio is not negligible but the slow-fast condition between variables and parameters still applies.

For this class, normal forms can be used to analytically study systems’ robustness and derive early warning signals for impending tipping points (Kuehn, 2011). Normal forms are general and low-dimensional models 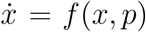 that describe topologically equivalent systems within a bifurcation class, in the vicinity of critical points (Kuznetsov, 2013). They allow to extract analytical and generic results for wide classes of systems (Kuehn and Bick, 2021), at the price of neglecting homeostatic dynamics far from tipping points. As a result, they allow to focus on critical transition mechanisms across various systems, instead of studying the full evolution of a single system. Details about topological equivalence and construction of normal forms are in STAR Method C.1. Fig. 2 shows an example of reduction to normal forms for two simple models.

**Figure 2:**
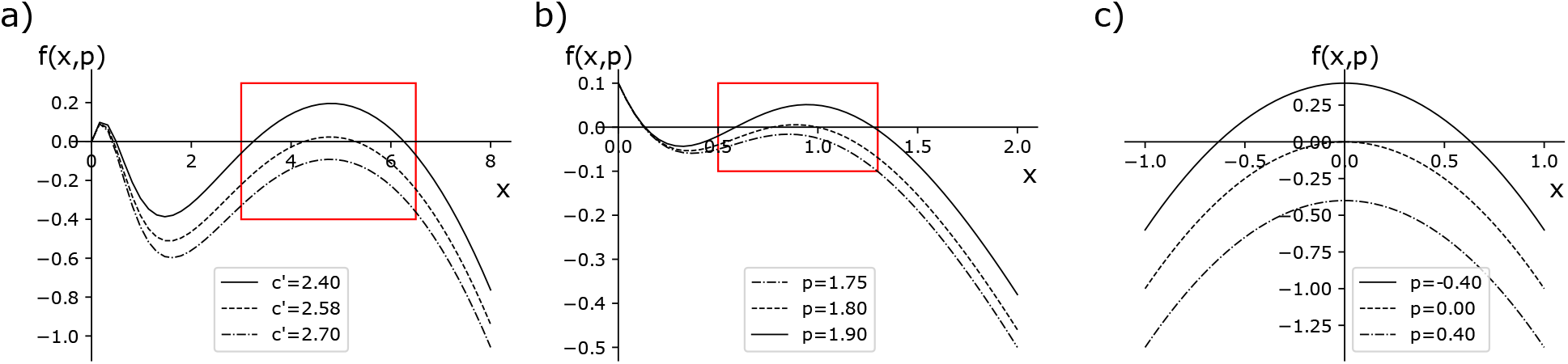
Visual example of topological equivalence. a) Plot for *dX/dt* = *f* (*x, p* = *c*′) = *X* (1 − *X/K*) – *c*′*X*^2^*/*(*X*^2^ + 1), a model of harvested ecological populations (Scheffer et al., 2009), also akin to Allee effects observed in microbiological colonies (Dai et al., 2012); *X* is the population density, *K* is the carrying capacity and *c*′ is the maximum harvest rate. b) Plot for *f* (*x, c*) of the autocatalytic loop model Eq. 15. c) Plot for *f* (*x, p*) of the saddle node normal form 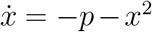. The two realistic models are locally topologically equivalent to the normal form within the red rectangle (visual reference): they approach a bifurcation point, marked by *f* (*x, p*) crossing the x-axis, as the parameter *c*′ or *c* changes.

Here, we consider those normal forms of primary biological interest. The saddle-node bifurcation, often associated with population collapses (Scheffer et al., 2009; Dai et al., 2012) or biological state transitions (Alon, 2006), is defined by *f* (*x, p*) = *± p ± x*^2^. At *p* = 0, a stable 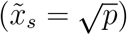 and unstable 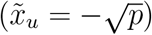 branch collide and vanish, resulting in a critical transition to an alternative branch (if it exists). Transcritical bifurcations *f* (*x, p*) = *px* − *x*^2^ are characteristic, for instance, of epidemic outbreaks (Proverbio et al., 2022a). Here, the two equilibria *x*_1_ = 0 and *x*_2_ = *p* meet at *p* = 0 and exchange stability. Finally, the family of pitchfork bifurcations *f* (*x, p*) = *px* + *l x*^3^ describe branching processes from one to two states (or viceversa); *l >* 0 identifies subcritical bifurcations, associated to critical transitions, while *l <* 0 defines the supercritical case, with a continuous transition over mean values. This mechanism is identified in cell regulation processes (Moris et al., 2016). Stochastically forced systems, associated to “noisy b-tipping”, can be written in the Itô form (Thompson and Sieber, 2011)

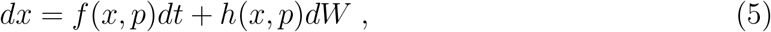

where *dW* is a Wiener process with variance *σ* and *f* (*x, p*) is a suitable normal form from those described above. The term *h*(*x, p*) allows to represent different noise types, to reflect modern knowledge of stochastic processes occurring in biological systems. Additive Gaussian noise with *h*(*x, p*) = 1 is usually associated to extrinsic cell-cell variability. State-dependent (multiplicative) noise *h*(*x, p*) ≠ const represents intrinsic stochasticity determined by, e.g., reaction rates, timescales or species concentrations of the underlying biochemical processes (O’Regan and Burton, 2018). Combinations of additive and multiplicative noise, with various ratios depending on different systems, are more realistic (Liu et al., 2009; Sidney et al., 2010) and fit experimental data better than Gaussian noise (Wang et al., 2012b).

If the microscopic kinetics is known, the noise terms can be exactly derived from the Master equation using Gillespie formalism (Gillespie, 2000). Alternatively, a diffusion approximation (Allen, 2010; Van Kampen, 1992) derives noise terms proportional to system state (*h*(*x, p*) = *x*), or to the drift term of Eq. 5, *h*(*x, p*) ∝ *f* (*x, p*). Here, for multiplicative noise, we consider 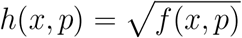 (O’Regan and Burton, 2018) and *h*(*x, p*) = *f* (*x, p*), to reflect modelling of biological regulatory circuits (Hasty et al., 2000). This way, mechanistic and stochastic normal-form bifurcation models are examined to study the effects of intrinsic and extrinsic noise on statistical patterns of variability and related EWS.

Following the procedure detailed in STAR Method C.2, Eq. 5 is analysed by solving the slow dynamics, linearising around a trajectory inside the stable (attracting) manifold and changing the coordinates to highlight the residuals *y*(*t*) around the linearization. This procedure gives

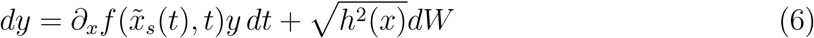

where 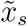 corresponds to the attracting part of the critical manifold (stable solutions). The linearised drift term corresponds to the leading eigenvalue of the deterministic normal form. Its magnitude 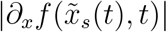 is the asymptotic decay rate of a perturbation. It corresponds to the concept of engineering resilience (Holling, 1996), which is akin to that of robustness (Kitano, 2004). A change of notation 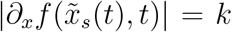 makes explicit that Eq. 6 corresponds to a (possibly non-autonomous) Ornstein-Uhlenbeck process, with critical *k* given by *k*_0_ = 0. It is a well-studied problem in stochastic processes theory, with analytical solutions for its statistics in different regimes (Allen, 2010; Gardiner, 1985). Eq. 6 can be regarded as a first order autoregressive model. However, its derivation from normal forms allows more nuanced interpretation: rather than being hypothesised as a statistical model to capture simple relationships, it is general for all models that can be reduced to normal forms.

## 2. Results

### 2.1 Robustness of EWS for noisy bifurcations

Within the class of critical transitions induced by bifurcations characterised by small fluctuations, discussed in Box 1 and 2, we study the early warning signals associated to impending tipping points, considering different noise types that are better representative of biological dynamics that pure Gaussian noise (see Box 2).

Analytic expressions for key summary statistics indicators can be obtained from Eq. 6 using standard approaches for stochastic processes (Allen, 2010; Gardiner, 1985). Their behaviour as the control parameter changes provides early warning signals for approaching noisy bifurcations (Scheffer et al., 2009). The lag-*τ* autocorrelation function does not depend on *h*^2^(*x, p*) but only on 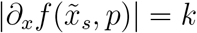:

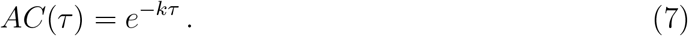

Hence, the common indicator lag-1 autocorrelation (AC(1), with *τ* = 1) only depends on the dampening rate. The power spectrum of the Fourier transforms and the variance, two common indicators, explicitly depend on *h*^2^(*x, p*):

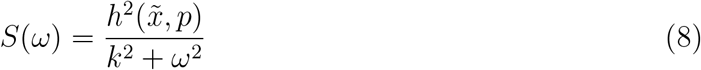

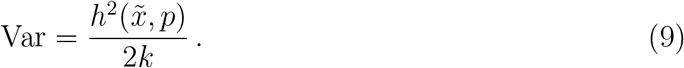

Coefficient of variation (CV) and Index of dispersion (ID), defined as

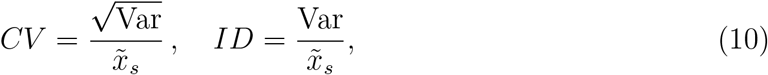

also depend on 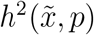. Other statistical moments, for stochastic processes with quasisteady state parameter, can be expressed as

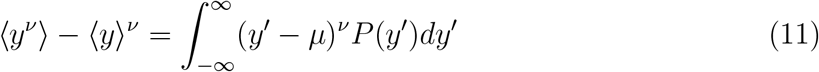

where *P* (*y*′) is the probability density function from the associated Fokker-Plank equation (Gardiner, 1985) and *μ* is the expected average value. Skewness and kurtosis, sometimes suggested as indicators for EWS (Guttal and Jayaprakash, 2008), can be easily extracted from Eq. 11 as third and fourth moments (*ν* = 3 and 4). Entropy-based indicators are more challenging to derive in case of multiplicative noise, as their defining integrals may not be solvable. Their derivation in case of Gaussian noise is described in STAR Method C.4; for the other cases, their behaviour is estimated below using computer simulations.

In all cases, the analytical results for each normal form can be obtained by substituting the corresponding dependency of the drift term to the control parameter: for the saddlenode, 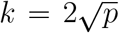, for the transcritical *k* = *p* and for the pitchforks *k* = 2*p*. In Fig. 3, the effect of multiplicative noise on the trends of common indicators is shown using 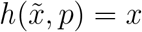 and 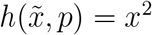.

**Figure 3:**
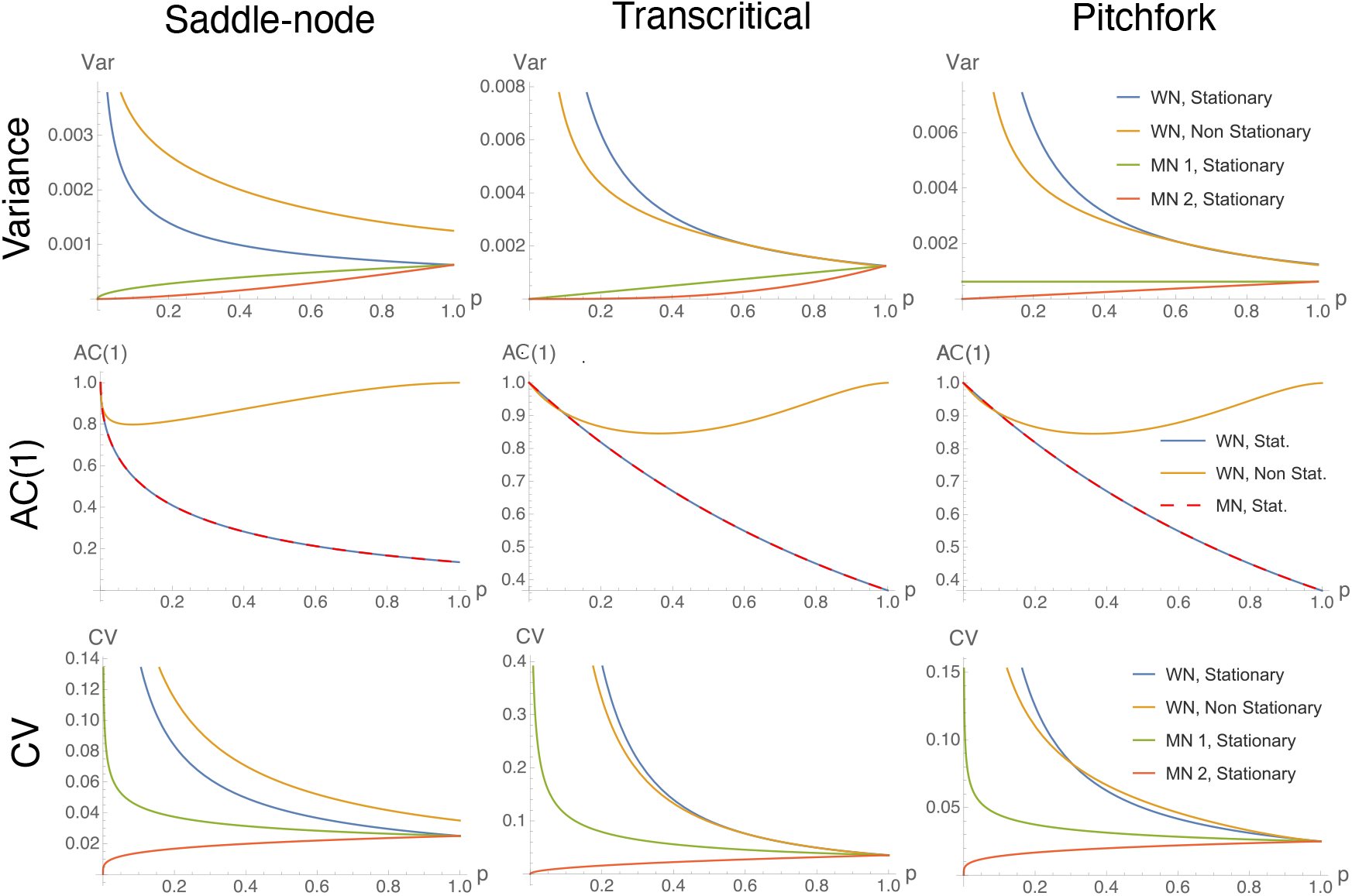
Trends of common statistical indicators. We consider Var, AC(1) and CV for saddle-node, transcritical and pitchfork bifurcations as *p* → 0, in different dynamical contexts (combinations of noise characteristics and stationarity for the control parameter). WN: white (Gaussian) noise; MN 1: multiplicative (state-dependent) noise 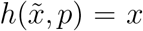; MN 2: multiplicative noise 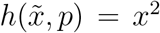. As the autocorrelation is independent on noise, only MN 1 is show and it overlaps with the white noise case.

Fig. 3 shows expected trends of common statistical indicators, for the three main normal forms and different noise types. Although the scaling induced by *k*(*p*) differs, the qualitative trends are conserved across the bifurcations. This observation suggests genericity of EWS, but also difficulties to infer the existence of one or another bifurcation using statistical indicators alone. Other methods (e.g. Angeli et al. (2004)) are recommended to complement the inference.

For Gaussian noise, EWS are associated with increasing trends of statistical indicators (Dakos et al., 2015; Scheffer et al., 2009). However, multiplicative noise may alter or completely disrupt them (as also noted by O’Regan and Burton (2018)), resulting in no early warnings prior to tipping points. Eq. 8 shows that even power spectrum trends can be subject to alterations from expected patterns, potentially resulting in spurious signals.

A preliminary investigation on ramping parameters (Pavithran and Sujith, 2021) can also be conducted. In this case, *τ*_*x*_ ≃ *τ*_*p*_: the quasi-steady-state (stationary) assumption is relaxed, but r-tipping may not yet occur. Let us consider linear ramping as *k* = *k*_0_ − *at*, where *k*_0_ is any initial condition, *a* is a small rate coefficient and the ramping stops at the critical value *k* = 0. Both coefficients are set to 1 to represent commensurable time scales. Only Gaussian noise is considered. This is a particular case of inhomogeneous processes (Gardiner, 1985) for which statistical moment solutions exist in the form

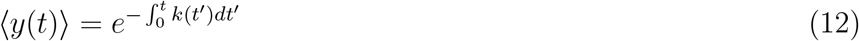

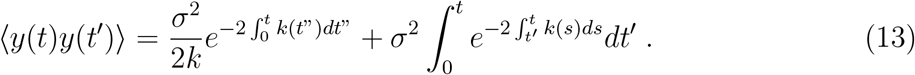

Derived statistics are calculated analogously. Eq. 13 is solved using Mathematica software to tackle the rightmost integral yielding the non-elementary Error function Erf(t). Fig. 3 shows that trends of common indicators may be modified by commensurable time scales of parameters evolution. Hence, raising reliable alerts becomes more challenging.

Overall, this analysis demonstrates that theoretical early warning signals due to increasing trends of summary statistics are sensitive to the “dynamical context”, i.e. noise properties and reciprocal time-scales. Hence, if the dynamical context is not carefully accounted for, spurious signals may be extracted from data, as observed in early findings from single systems (Brett et al., 2017; Proverbio et al., 2022a).

If the context is known, the current results suggest which indicators to use to obtain robust early warnings. The autocorrelation is robust against changing noise properties; the variance is more sensitive to multiplicative noise, but maintains its expected trends in case of ramping parameters. The coefficient of variation is also robust in case of commensurable time scales and copes well in case of certain types of multiplicative noise. Overall, what matters is the competition between changes in noise and changes in resilience: depending on which one is more rapid, the indicators and their associated EWS may perform as expected or fail to anticipate an impending critical transition.

Measurement processes or details of realistic models may further influence EWS. Measurement uncertainties, assumed as Poisson processes associated with measuring instruments or procedures and thus independent of systems’ dynamics, can be introduced in the formulas of statistical indicators by error propagation in quadrature (see STAR Method C.4 for details). In case of Gaussian noise and stationary processes, the expected trends of common indicators are not altered, hence, EWS can be in principle extracted even when using noisy measurements (*cf*. STAR Method C.4).

Single indicators may also be skewed in case realistic details are considered. For instance, on empirical data, normalising by the critical value and set a normal form around *p*_0_ = 0 and 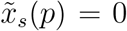 may be challenging, since such critical values are largely unknown. Hence, instead of computing 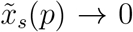 like on perfectly reconstructed normal forms, 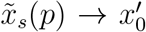 is often computed (Dai et al., 2015), where 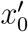 corresponds to the critical value, unknown a priori. Such case can be modelled as 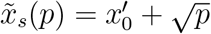. Hence, Eq. 10 becomes

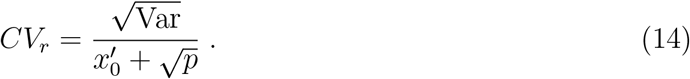

Here, other multiplicative noise forms may alter its behaviour and shadow possible early warnings. Finally, skewness and kurtosis calculated from Eq. 11 display increasing trends when *P* (*y*′) is symmetric (STAR Method C.4). However, this may not be true in case of multiplicative noise (Sharma et al., 2016), resulting in distorted trends and early warnings. In this sense, there is no ambiguity between the results of Guttal and Jayaprakash (2008), proposing EWS from skewness, and Dai et al. (2012), observing flat and fluctuating trends on experimental data: likely, the noise properties were different than what assumed.

### 2.2 Optimisation of EWS

Having assessed in which cases the proposed early warning signals are expected to work for noisy b-tipping transitions, we now optimise their performance to provide significant and as-early-as-possible alerts, in a range of dynamical contexts and for the most common transitions observed in systems biology. To this end, we focus on multistable systems (Sarkar et al., 2019), develop and solve an optimisation problem using computer simulations to go beyond the first-order approximation from Eq. 6 (see STAR Method C.2 for details), and study a wide range of noise levels and types, to establish a composite indicator that is robust and performing across multiple systems.

Multistable systems are systems whose deterministic landscape features at least two attractors (Feng et al., 2016), and usually undergo either saddle-node bifurcations or ntipping. Bistability means local multistability across two attractors. Angeli et al. (2004) provides necessary and sufficient conditions for bistability in a wide range of biological systems. Among them, a feedback model with three-points I/O characteristic curves suffices. A simple linear system with monotonic sigmoidal feedback can do the job, in a range of parameters (Fig. 4). As a case study, the autocatalytic positive feedback loop derived from Michelis-Menten kynetics (Sharma et al., 2016)

**Figure 4:**
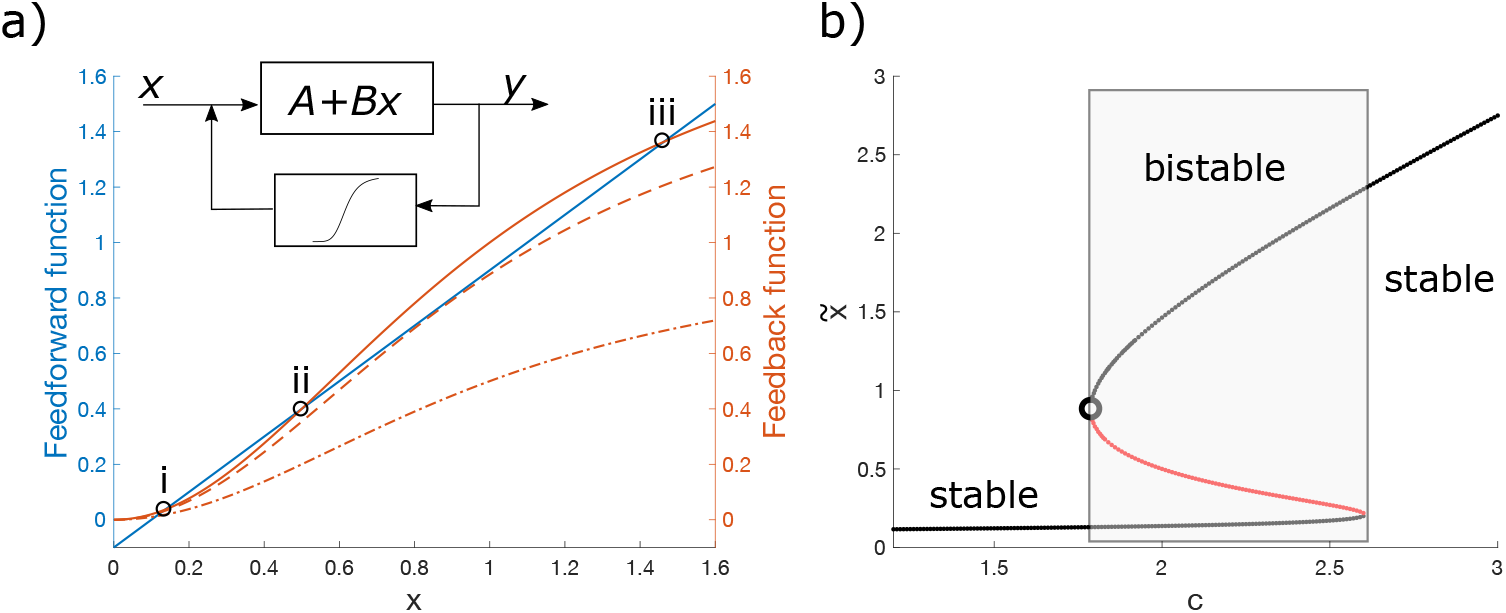
Bistable systems, studied with (a) characteristic curves or (b) bifurcation diagram 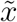 (stable state) vs control parameter. Among the systems undergoing saddle-node bifurcations, any linear system with nonlinear feedback and adequate feedback gain, such that the characteristic curve crosses the activation function in three points (two stable, one unstable), can display bistability. This example uses Eq. 15. a) The feedback function *FB* corresponds to the Hill function (*k* = 2), the feedforward *FF* to the linear part −(*K* − *x*) with *K* in its appropriate range. The control parameter *c* tunes the *FB* function. Dashed-dotted line: *c* is not sufficient to promote bistability, corresponding to left stable region of (b). Dashed line: the critical value for which *FB* is tangent to *FF*, corresponding to saddle-node point, open circle in (b). Solid line: bistable system with three intersection points (stable, i and iii; unstable, ii). When studying the vector field *f* (*x, c*) is easier than the characteristic curves, one can use the representation and interpretation in Fig. 2b. Note that the line styles have the same meaning in panel (a) and Fig. 2.

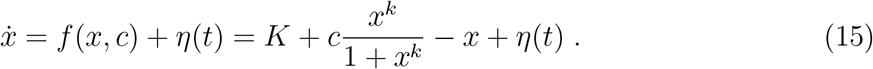

satisfies the bistability conditions, and can thus display transitions between attractors, if 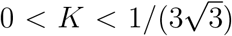 for *k* = 2 (Weber and Buceta, 2013). In Eq. 15, *x* is the concentration of a transcriptional factor activator, activating its own transcriptions when bound to a responsive element; *K* is the basal expression rate, *c* is the maximum production rate, *k* is the Hill coefficient and *η*(*t*) accounts for the stochastic terms. Eq. 15 comes from a two-variable genetic toggle switch, assuming slow-fast timescale separation between the two variables (Strogatz, 2015) and after a-dimensionalising the chemical details to retain the dynamical scaffold. Notably, networks of Michelis-Menten regulators can be reduced to Eq. 15 after dimension reduction techniques (Gao et al., 2016).

Eq. 15 displays bistability for a range of values *c* (the exact range depends on *K* and *k* (Proverbio et al., 2022b)) and, in particular, a saddle-node bifurcation between two alternative steady states at a critical value *c*_0_ of the parameter *c*, such that 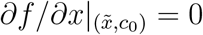:

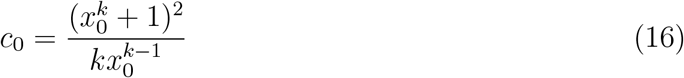

where *x*_0_ is the tipping value for the system state. Therefore, system 15 can be used as a paradigmatic example of biological systems, within the saddle-node b-tipping class, to perform optimisation studies that go beyond the local and low-noise-to-signal-ratio approximation provided by normal forms.

The quasi-steady state assumption is generally accepted for such systems (Del Vecchio et al., 2016), so we focus on dynamical contexts characterised by different types of noise, whether yielding n-tipping or possibly skewing statistical indicators due to multiplicative and/or additive nature. To model combinations of intrinsic and extrinsic noise, we set

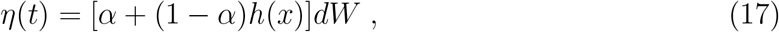

where *α* weights the white or multiplicative noise component (*α* = 1 corresponds to purely additive Gaussian noise, *α* = 0 to purely multiplicative); like above, *h*(*x*) = *x* or *h*(*x*) ∝ *f* (*x*) = *x*^*k*^*/*(1 + *x*^*k*^) (Hasty et al., 2000) and *dW* is a Wiener process with variance *σ*. Without loss of generality (Proverbio et al., 2022b), we set *k* = 2.

As early warning signals are associated with increases of statistical indicators, we need to establish a measure of statistically significant increase, to rule out false positives and false negatives due to random fluctuations in the indicators. To do so, we employ the p-value analysis used in Proverbio et al. (2022b) (see STAR Method C.6 for details). It allows to measure at which value of the control parameter *c*, before *c*_0_, a significant signal is triggered, thus obtaining a “lead-parameter” 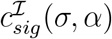 depending on noise properties and the considered indicator ℐ (see STAR Method C.6 for details). 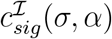 is first computed for each indicator individually. Fig. 5a shows the results in case of white noise, while various functionals of multiplicative noise *h*(*x*) (with *α* = 0) are reported in Supplementary Figure S3. Each indicator yields various *c*_*sig*_; in Fig. 5a, Var, AC(1) and *H*_*S*_ maximise *c*_*sig*_ over various noise levels, while other indicators like skewness and kurtosis perform poorly, as anticipated by the analytical results. CV and ID are also rather poor, likely due to fluctuations of mean values and anticipating n-tipping (*cf*. also Supplementary Figure S2 and S3). For the case of multiplicative noise (Supplementary Figure S3), *H*_*S*_ keeps performing well while Var, as expected from the theoretical analysis, decreases its performance despite being still better that Skew and Kurt.

**Figure 5:**
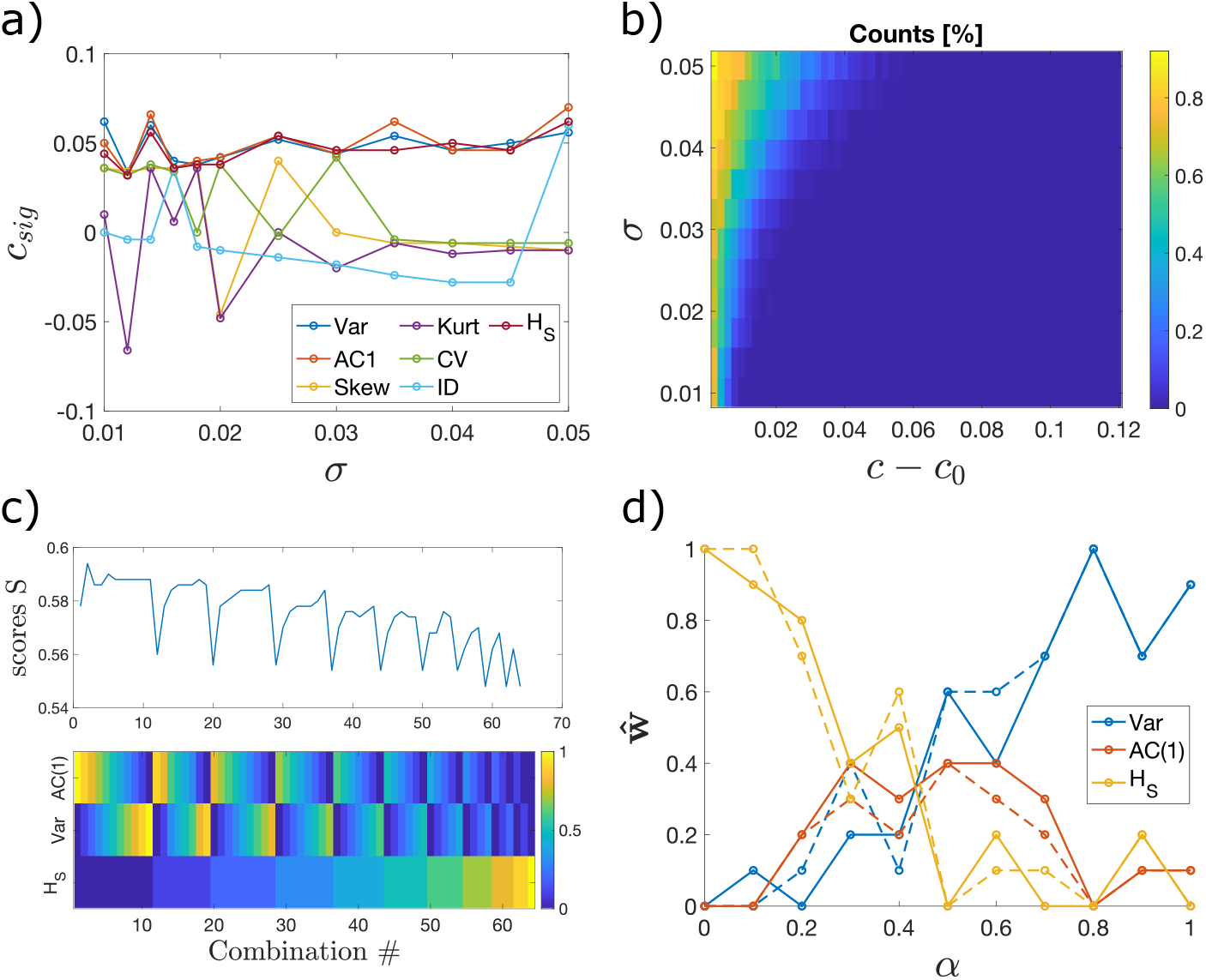
Optimisation of leading indicators for EWS, according to lead parameter *c*_*sig*_. a) *c*_*sig*_ at various noise intensities *σ*, for all the most common indicators. b) The counter 𝒞, normalised by all transitions to be interpreted as probability of n-tipping, at different noise intensities *σ* and distances *c* − *c*_0_ from the bifurcation point. c) Scores S, corresponding to the argument of the cost function Eq. 19, for various combinations 𝒮 from Eq. 18. In the panel below, the color code shows the weights *w*_*k*_ for each indicator, in each combination. Results in panels a, b and c refer to *α* = 1. For various types of *h*(*x*) and *α* = 0, see Supplementary Figure S3. d) Optimal weights **ŵ** for each indicator, as a function of noise mixing *α*. As a representative of multiplicative noise, we used *h*(*x*) = *x*. Other *h*(*x*) conserve the trends, albeit changing the corresponding *c*_*sig*_. It may happen that the optimisation is solved by multiple combinations (dashed lines).

Complementing the analysis of the lead parameter requires understanding how many noise-induced tipping events occurred before it and assessing whether the increasing indicators alert for impending collapses or reflect transitions that have already happened. The analysis thus interprets warning indicators as “anticipating” or “just-on-time detecting” the tipping events. To do so, a counter 𝒞 quantifies, for each parameter value *c* and for each noise level *σ*, how many trajectories tip onto the alternative stable state. The results are in Fig. 5b: as *σ* increases, more n-tipping events occur before the bifurcation point. In particular for *σ >* 0.42, several noise-induced transitions occur at *c* ≃ *c*_*sig*_. Hence, as noise increases, the indicators capture ongoing critical transitions but are not able anymore to provide much earlier alerts. This likely explains the remarks from Dudney and Suding (2020), that EWS could not anticipate several transitions in real-world systems, in particular those characterised by high noise-to-signal ratios.

The previous results are also employed to define an optimisation problem to maximise 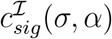 for varying *α*. To do so, we define a composite indicator as linear combination of indicators

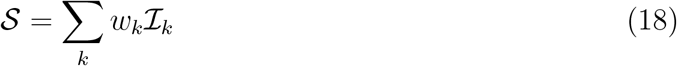

and look for a set of weights **w** = {*w*_*k*_} that maximises all 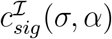 as *σ* increases (to guarantee robustness against noise levels), for the various *α*:

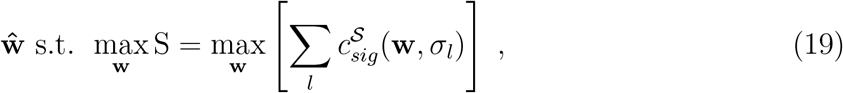

where S are scores composed by sums of 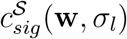 over all *σ*. In the set ℐ, we include those indicators that are expected to be robust and performing, first and foremost in the white noise case. Leveraging on the previous results, we therefore select Var, AC(1) and *H*_*S*_. As the problem is non-convex (Fig. 5c), we perform a grid search for all combinations of *w*_*k*_, with a stride 0.1 and such that ∑_*k*_ *w*_*k*_ = 1. See Fig. Fig. 5c for the considered combinations to construct 𝒮.

Figure 5d reports the results of the optimisation procedure. Combinations of Var and AC(1) make up for optimal indicators in case of white noise, **ŵ** = [0.9, 0.1, 0] for Var, AC(1) and *H*_*S*_, respectively, in case of *α* = 0. In this case, *H*_*S*_ is log-proportional to Var (see Eq. C.11) and does not add much information. In turn, combining the indicators maximises 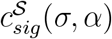 in case of mixed noise types. Finally, when multiplicative noise is prevalent in the system, using Shannon entropy is preferred (**ŵ** = [0, 0, 1] for *α* = 0). Note that, as the problem is non-convex, there may be more than one combination to create the optimal 𝒮. However, changes in weights *w*_*k*_ are always within Δ*w*_*k*_ ∼ *±*10%*w*_*k*_ and the trends are conserved (see dashed lines in Fig. 5d). Such small Δ*w*_*k*_ yield changes of *±*4% on the scores S, on average over all *α* (ΔS ∈ [1.8; 6.5]%), while off-setting *w*_*k*_ by more than 50% (e.g., using full variance in case of multiplicative noise) worsens S (and consequently the optimal lead parameter) up to more than 20%.

### 2.3 Verification on experimental data

The theoretical predictions are verified and used to interpret experimental data from a previous publication (Dai et al., 2012). The data are sampled from controlled experiments of budding yeast population collapse. Budding yeast cooperatively breaks down the sucrose necessary for its survival, thus inducing a density-dependent dynamics that realises the Allee effect of bistable population dynamics (*cf*. Fig. 2b). Repeated experiments empirically reproduced a saddle-node bifurcation by measuring population density (state variable) as a function of dilution factors (DF, control parameters) affecting the sucrose environment. Various EWS for population collapse can be estimated using distributional data. More details about data collection and analysis are in STAR Method C.7. Testing our theoretical results on a different system than Eq. 15, yet still belonging to the saddle-node driven b-tipping class, would thus assess their generic applicability within this class.

Fig. 6 shows trends of each indicator individually, as function of the dilution factor (with critical value at 1600). The error bars are estimated from bootstrapping (STAR Method C.7). Fig. 6 reproduces the results from Dai et al. (2012) and includes the additional indicators considered in this paper. The mean is used to reconstruct the upper stable branch of a saddle-node bifurcation diagram (see Fig. 1), reconstructed from data (the full diagram can be found in the original publication). However, it can not be used as proper EWS as decreasing mean values could signify smooth changes rather than critical transitions, if the transition type and critical parameter are not known. Skewness and kurtosis fluctuate around 0 and 3, respectively, without providing EWS, as one expects in case of symmetric potentials (see Eq. C.17 and C.18). AC(1) and the autocorrelation time (defined as − 1*/* log[*AC*(1)] (Dai et al., 2012)) first drop before increasing sharply just before the critical value. Comparing it with Fig. 3, we speculate that there are commensurable time scales between the intake of sugar by yeast cells and their evolution in density. Further experiments are suggested to check for this intriguing hypothesis.

**Figure 6:**
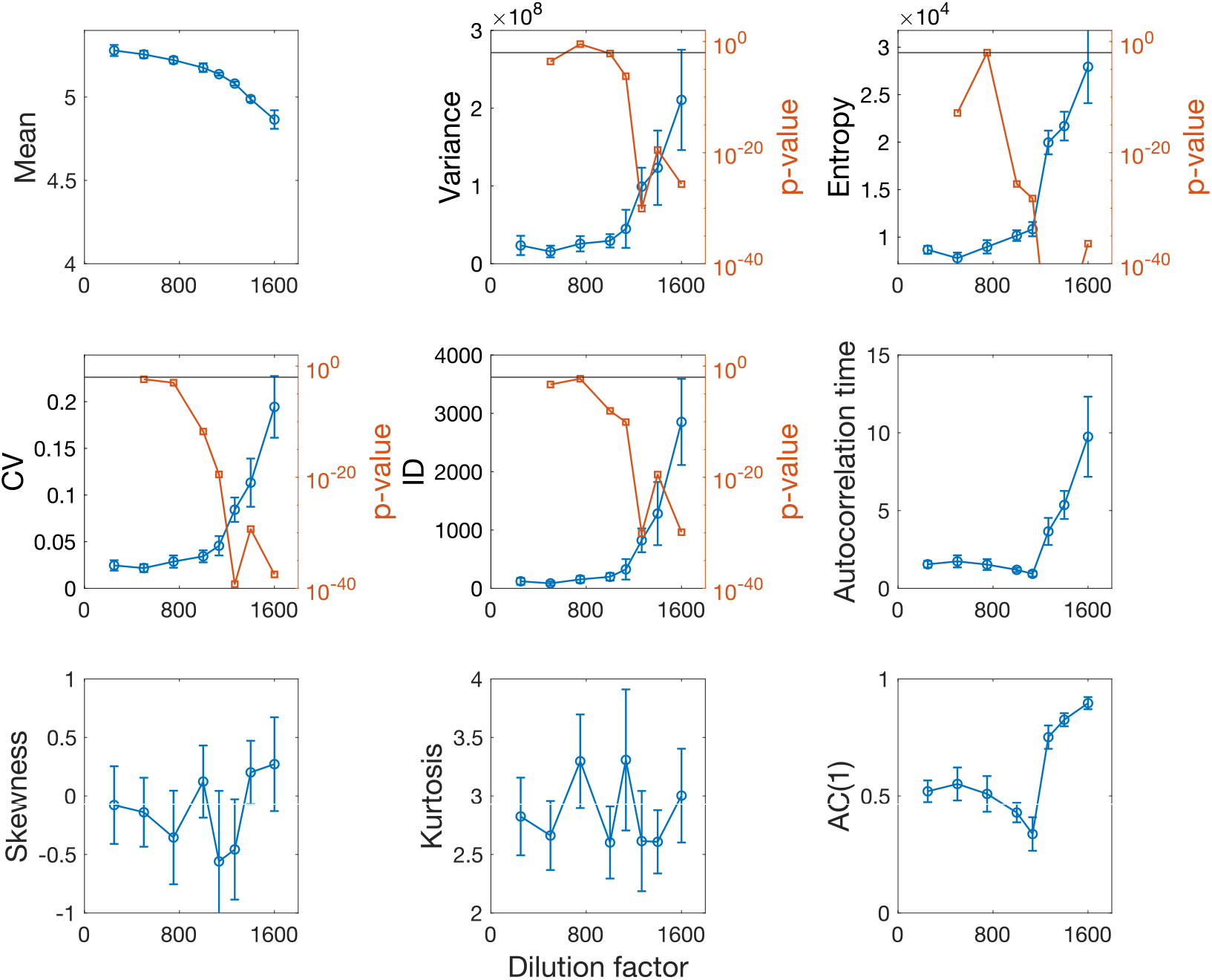
Statistical indicators calculated on empirical data. Data are from Dai et al. (2012), as functions of dilution factor (DF). Their corresponding p-values are estimated when the trend is increasing while approaching the bifurcation point (rightwards point). All statistical moments of degree *γ* have units of measure (cells/*μl*)^*γ*^. The autocorrelation time is in days. The mean reproduces the upper stable branch of a saddle-node bifurcation diagram (*cf*. Fig. 1) until the empirically estimated bifurcation point at DF=1600. Horizontal solid lines mark p-value = 0.01.

Even in this case, as expected, Var, Entropy (*H*_*S*_), CV and ID display monotonous increasing trends close to the bifurcation point. The increases are thus assessed using the p-value test (*cf*. STAR Method C.6) to check whether they are significant or associated with fluctuations. To trigger a significant early warning signal, we require a conservatory significant p-value*<* 0.01. This way, we estimate the significant dilution factor DF_*sig*_ for each indicator. For variance, DF_*sig*_ = 1133, for the others DF_*sig*_ = 1000. Comparing with the optimisation results (from the previous section and Fig. S3), we infer the presence of multiplicative noise in the system’s dynamics. Note that entropy showcases the smallest p-value at DF_*sig*_ = 1000; it is also the most robust when changing the repetitions in the bootstrapping procedure (STAR Method C.7).

To test the hypothesis of association between EWS performance and noise type, we test combined indicators with *H*_*S*_ and Var. According to the optimisation above, the higher the variance content in the mixture, the lower the significance of the increasing trend. This is what is observed in Fig. 7: having a balance between Var and *H*_*S*_ yields DF_*sig*_ = 1000, but with a higher p-value than when reducing the ratio Var/*H*_*S*_ or when comparing with the case of entropy alone (from Fig. 6).

**Figure 7:**
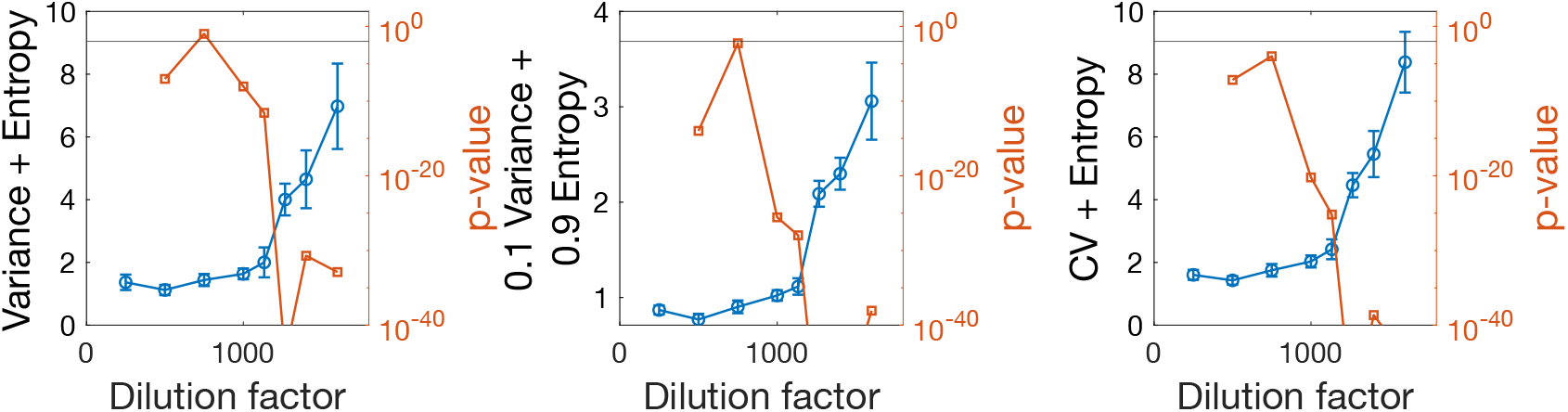
Combined indicators calculated on empirical data. The analysis is analogous to that in Fig. 6, using combinations of indicators. The horizontal solid lines mark p-value = 0.01.

Finally, we test combining CV and *H*_*S*_, since Fig. 6 suggests that CV could perform well. Indeed the new combined indicator yields DF_*sig*_ = 750 (Fig. 7, right), one dilution step before the others. This is not in contrast with the optimisation analysis: CV is, in fact, expected to be as performing as *H*_*S*_ if the noise levels are relatively high (see Fig. S3). We recall that CV was not included in the optimisation analysis to be generic and robust across noise types and levels. However, if high *σ* in state-dependent noise is known, constructing a composite indicator using both CV and *H*_*S*_ may improve the alerting performance.

## 3. Discussion

The paper provides a systematic classification of tipping mechanisms, highlights their underlying modelling assumptions, and bridges mathematical insights and observations of real systems to classify various tipping mechanisms, towards quantitative understanding and prediction of such relevant phenomena. The work shifts the focus from studying specific systems, that may undergo some transitions, to studying transitions, along with their classes and properties, which can accommodate the behaviour of different systems. An interesting question for future studies will be to develop data-driven methods to classify each system within its corresponding class, much like those developed to distinguish stochastic or chaotic signals (Rosso et al., 2007). This will dramatically help the understanding of biological processes and guide the selection of EWS or other methods to anticipate critical transitions, as well as informing methods to reconstruct cell developmental trajectories like those proposed by Eugenio et al. (2014).

Moreover, we systematically investigate early warning signals associated to noisy bifurcation-induced transitions, key dynamical routes for the regulation and control of many natural processes. So far, EWS have been mostly studied in highly controlled computational settings, or checked on empirical data with alternate success. Our results make sense of previous observations, help to define their range of applicability to reliably predict systems’ behaviours, and allow to understand why spurious signals may be triggered in certain cases. We also assess whether and when EWS can be interpreted as anticipating or just-on-time detecting critical transitions in the presence of noise. By carefully analysing noise types and parameter dynamics, we also extend previous results to more realistic settings, to guide real-world applications.

Using both analytical and computational methods, we observe that the variance – a highly employed indicator for EWS – may be sensitive to state-dependent noise, while AC(1) can be skewed by ramping control parameters. Both are good indicators in case of quasi-steady-state dynamics and Gaussian noise, with the ability to provide information about augmented risk of tipping events. In the other cases, Shannon entropy is the most robust and performing indicator and is suggested for applications in case of uncertain settings. If precise information about noise type and intensity are available, constructing composite indicators can improve the early-alerting performance, e.g., by combining CV and *H*_*S*_.

The optimisation of composite indicators points to the use of machine learning methods when abundant data are available (Bury et al., 2021), but also opens important caveats for their application in real life: feature combinations may be optimised for certain settings (e.g., noise intensity or type) but may be hardly generalisable for others. Our results remark that training should be performed considering all possible combinations, or by first assessing which critical transition class is being considering. Otherwise, misleading signals may be triggered and wrong conclusions reached. On the other hand, our results can be used for feature selection of more interpreteable machine learning algorithms that leverage the proposed composite indicators, insofar defined for a-priori assessment of systems that lack big data.

This work provides results and guidelines for the application of early warning signals from the critical transitions framework, but some points should still be covered by future studies. They include more refined analytical derivations of indicators in case of inhomogeneous processes as well as closed formulae for entropy in exotic settings. Further investigations on realistic systems, including non-autonomous transitions currently understudied in systems biology, are thus suggested as extensions of our work. Another limitation of the present study is the restriction to low dimensional systems. In principle, they are representative of any system after dimension reduction techniques are applied, but it is necessary to assess if and how the latter induce performance drops. Extending the analysis to high dimensional systems, e.g., by testing multivariate indicators (Weinans et al., 2021) or further refining EWS performance when multiple independent variables can be observed, is thus suggested to future studies. Finally, our theoretical results have been verified on empirical data from literature, but we acknowledge the need of performing additional experiments to continuously validate our predictions. In particular, we suggest to design new experiments to test the quantitative predictions about lead parameters and to assess what happens in case of rapidly ramping parameters.

Our results can be readily tested and applied on real-world monitoring systems and can inform the development of new indicators to address specific problems like cancer onset, much like previous works (Chen et al., 2012) did using less performing measurements. In addition, leveraging the sensitivity of indicators’ trends to noise type and parameter dynamics can provide new methods to infer the latter from empirical data. For instance, comparing Fig. 6 with Fig. 3 supports hypothesis of commensurable time scales between intake of sucrose (affected by the dilution factor) and cells’ growth in yeast experiments (Dai et al., 2012); such hypothesis, to be confirmed using controlled experiments, could advance our knowledge beyond the current slow-fast approximations (Del Vecchio et al., 2016). Similarly, the prevalence of certain noise types can be inferred by comparing data and theory. Overall, we connect theory and data, such that knowledge about the dynamical settings allows optimising early warning signals, and analysis of statistical indicators enables inference of dynamical properties.

## Acknowledgement

The authors would thank their colleagues for valuable discussions. D.P.’s work is supported by the FNR PRIDE DTU CriTiCS, ref 10907093, and A.S. by the FNR (C14/ BM/ 7975668/ CaSCAD) and by the NIH NBCR (NIH P41 GM103426).

## Author contributions

Conceptualization, D.P and A.S. and J.G.; methodology and investigation, D.P.; manuscript writing, D.P and A.S. and J.G.; supervision, A.S. and J.G.; funding acquisition, A.S. and J.G.

## STAR Method A. Key Resources

Software: Matlab R2021b (Matworks), Mathematica v12 (Wolfram)

Analysis and figures script: GitHub (https://github.com/daniele-proverbio/EWS_optimise)

## STAR Method B. Resource availability

### STAR Method B.1. Material availability

This study did not generate new materials

### STAR Method B.2. Data and code availability

- All original code has been deposited at GitHub, https://github.com/daniele-proverbio/EWS_optimise, and is publicly available.
- All data used are publicly available on Zenodo: Dai, L., Vorselen, D., Korolev, K. S., Gore, J. (2012). Generic Indicators for Loss of Resilience Before a Tipping Point Leading to Population Collapse. Science. https://doi.org/10.1126/science.1219805

## STAR Method C. Method details

### STAR Method C.1. Topological equivalence and normal forms

Bifurcations model drastic changes in the qualitative behaviour of dynamical systems, such as shifts in equilibria and regimes (Kuznetsov, 2013; Kuehn and Bick, 2021). Before delving into bifurcations and their representation as normal forms, recall the concept of topological equivalence.

Local topological equivalence between two dynamical systems {𝒯, ℝ^*n*^, *ϕ*^*t*^} and {𝒯, ℝ, *ψ*^*t*^} is established if there exist a homeomorphism *h* : ℝ^*n*^ → ℝ^*n*^ that maps orbits of the first system to orbits of the second one, and the direction of time is preserved. Local topologically equivalence near an equilibrium *û* is, in turn, established between a dynamical system {𝒯, ℝ^*n*^, *ϕ*^*t*^} and a dynamical system {𝒯, ℝ, *ψ*^*t*^} near an equilibrium *ŷ* if there exist a homeomorphism *h* : ℝ^*n*^ →ℝ^*n*^ that is defined in a small neighborhood *U*∈ ℝ^*n*^ of *û*, satisfies *ŷ* = *h*(*û*), and maps orbits of the {𝒯, ℝ^*n*^, *ϕ*^*t*^} ∈ *U* onto orbits of {𝒯, ℝ, *ψ*^*t*^} ∈ *V* = *h*(*U*) ⊂ ℝ^*n*^ while preserving the direction of time.

A bifurcation consists in the appearance of a topologically non-equivalent phase portrait under variation of parameters. The difference between the dimension of the parameter space and the dimension of the corresponding bifurcation boundary is called “codimension”. To determine a system’s behaviour near bifurcations, minimal-order forms, called “normal forms”, can be employed. In fact, the normal form of the bifurcation is locally topologically equivalent near an equilibrium to all systems exhibiting that certain type of bifurcation (Haragus and Iooss, 2010).

Consider a dynamical system

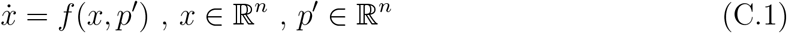

and a polynomial model

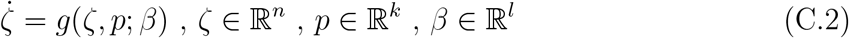

having dimension *n*, codimension *k* and polynomial order *l*. Without loss of generality, a change of coordinates can set the bifurcation point occurs at (*x, p*) = (0, *p*_0_) (Strogatz, 2015). System C.2 is thus called a *topological normal form* for a given bifurcation if any generic system C.1 with the equilibrium *x* = 0 satisfying the same bifurcation conditions at *p*′ = 0 is locally topologically equivalent near the origin to model (C.2) for some values of the coefficients *β*_*i*_. Using normal forms, it is thus possible to study classes of bifurcations using simple polynomials. If the system satisfies certain conditions on 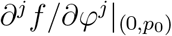 around the critical point, where *j* is the derivative order and *φ* = {*x, p*}, it is called “generic”. The nondegeneracy conditions ∂^*j*^*f/*∂*x*^*j*^ are related to the “criticality” of a bifurcation (Kuehn, 2011), while the trasversality conditions ∂^*j*^*f/*∂*p*^*j*^ govern the bifurcation unfolding and thus its genericity (the bifurcation exists even after small perturbations). The saddle-node investigated in the main text (*cf*. Fig. 4) is the most common generic normal form with dimension 1 and codimension 1 (Haragus and Iooss, 2010).

For low-dimensional systems, their associated normal forms can be explicitly obtained using e.g. Taylor expansion methods over both nondegeneracy and trasversality conditions (Strogatz, 2015). For high-dimensional systems, numerical methods like XPP-AUT (http://www.math.pitt.edu/~bard/xpp/whatis.html) or network reduction techniques (Gao et al., 2016; Tu et al., 2021) can be employed to infer or derive the normal forms. Obtaining analytical results for any system is still an open research field.

### STAR Method C.2. Analysis of slow dynamics

The fluctuations around the stable manifold of Eq. 5 can be analysed by studying the fast-slow dynamics around it and determining stochastic equations for the residuals (Kuehn, 2011; Berglund and Gentz, 2006; O’Regan and Burton, 2018). Here, we briefly recall the procedure to derive Eq. 6. Recall the normal form of a generic fold bifurcation:

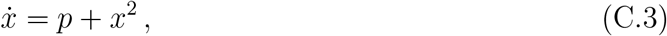

It has two steady states:

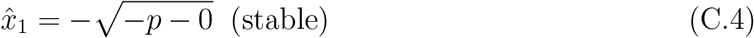

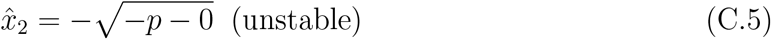

where the term “−0” explicits the distance from the bifurcation point *x*_0_ = 0 (by definition). Consider a neighborhood of the attractor (stable fixed point) 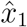 and see what happens after small perturbations. To do so, perform a local linearization by considering 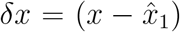. Thus:

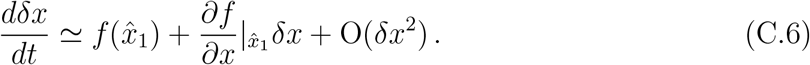

So, using Eq. C.3 and Eq. C.4, we obtain:

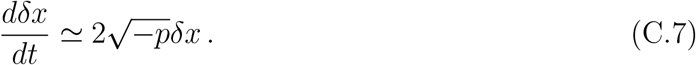

This deterministic form con be augmented by a Wiener process with variance *σ* arbitrary multiplied by *h*(*x*), representing non-Gaussian noise properties. This modelling choice converts the family of ODEs into SDEs (Berglund and Gentz, 2006; Namachchivaya and Leng, 1990; Khas’ minskii, 1966). A change *δx* → *y* makes the notation lighter into:

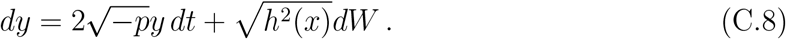

The equation describes a system evolving under small noise in a neighbourhood of the stable equilibrium, when this is not far away from the bifurcation point.

The term 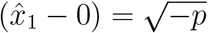 is the distance of the stable equilibrium from the bifurcation point and depends on the leading parameter *p*. We can thus rescale it to a new variable −*k*:

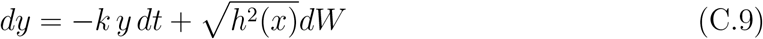

The sign “− ″ in “− *k*″ is included so that Eq. C.9 is interpreted as the associated Langevin equation to a Ornstein-Uhlenbeck process (Gardiner, 1985). The term multiplying the deterministic drift can thus be interpreted as −∂*V/*∂*x* where *V* (*x*) is the potential governing the drift of the particle subjected to random noise. In our case, thanks to the choices made,

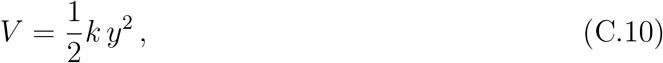

that is, a quadratically shaped adjoining potential typical of an overdamped oscillator under noise, of which *k* represents the depth. The working hypothesis is that boundary of the ideal potential *V* can grasp the boundary of the attracting basin of the original model after sufficiently long time. Eq. C.9 is analytically tractable to understand the main qualitative features of more complicated critical transitions. However, it requires ad hoc extensions when studying system-specific quantitative details like observability boundaries and lead times. Gardiner (1985) also extends Eq. C.9 to inhomogeneous processes with ramping parameters, used in Eq. 13.

### STAR Method C.3. Reproduce Fig 1

Fig. 1 displays examples of a bistable system with critical transitions and hysteresis as well as smooth transitions. Panel (a) corresponds to the bifurcation diagram of Eq. 15, flipped along the vertical axis to highlight the hysteresis.

Panel (c) shows the bifurcation diagram, over an unfolded supercritical pitchfork bifurcation, of 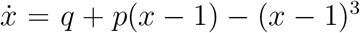, which corresponds to the bifurcation normal form, shifted (to better visualize the diagram) and modified by a small perturbing term *q* = 0.01 unfolding the bifurcation (Thompson and Sieber, 2011) into a smooth branch. In brief, an *unfolding* of a dynamical system under static equivalence is one that exhibits all possible bifurcations of the equilibrium (rest) points, up to topological equivalence of the set of equilibria (Kuznetsov, 2013). In other terms, it investigates what happens when small terms are added to the original bifurcation, mimicking extra parameters, small offsets or “impurities”. The illustrative attractors in panel (a) and (b) are two-well potentials associated, e.g., to the cusp bifurcation (aka “organising centre” Thompson and Sieber (2011); Eugenio et al. (2014)), a generic bifurcation described by 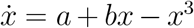, where the combination of *a* and *b* determine bistability and the route to a saddle-node bifurcation.

### STAR Method C.4. Supporting analytical results

#### STAR Method C.4.1. Entropy in case of Gaussian noise

Within a symmetric potential, elicited by a (locally) quadratic normal form, consider a Gaussian distributed variable *y* ∼ 𝒩 (*μ*, Var). Its entropy is:

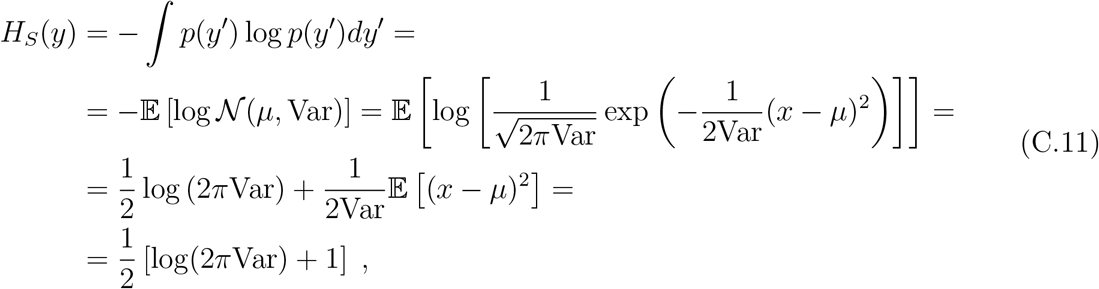

That is, for the case of Gaussian noise, *H*_*S*_ is directly proportional to the variance and displays similar trends, that can be used to derive EWS.

#### STAR Method C.4.2. Measurement noise

Consider a measurement process with uncertainties 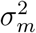, independent from system variance (Eq. 9). The resulting expected error, obtained from summing the two standard deviation in quadrature (Taylor, 1997), is:

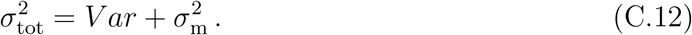

To derive the autocorrelation, combine its definition

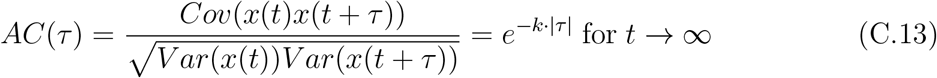

(where *Cov* indicates the covariance and *V ar* the variance) with Eq. C.12 (substituting 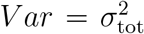). In principle, we can explicitly consider multiplicative noise like in the main text. However, the goal in this case is to compute if notable discrepancies exist between ideal measurements (no uncertainty) and realistic measurements (with some uncertainty, that can be filtered to correspond to white noise). Hence, only the case of white process noise is currently considered. This results in:

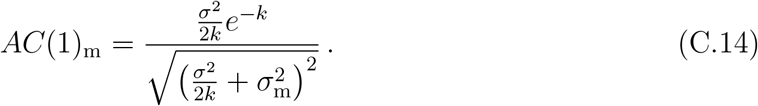

Obviously, 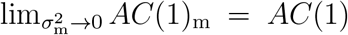. Fromm Eq. C.14, we can immediately see that measurement uncertainties *σ*_*m*_ induce small scaling but do not alter the functional. Only relatively high measurement uncertainty levels change the absolute values of expected lag-1 autocorrelation, but maintain the increasing patterns close to critical points.

#### STAR Method C.4.3. Skewness and kurtosis

For certain simulated systems, the third statistical moment (skewness) has been suggested to provide useful early warnings (Guttal and Jayaprakash, 2008). However, experimental results (Dai et al., 2012) were not able to confirm the expectations, estimating flat and fluctuating trends before a tipping point.

For a stochastic process with quasi-steady state parameter, its statistical moments are

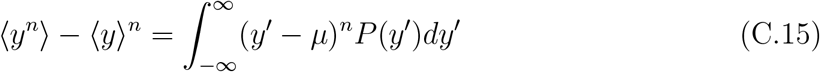

where *P* (*y*′) is the associated probability density function and *μ* is the expected average value.

For odd *n*, if *μ* = 0 and *P* (*y*′) is symmetric, the integral equals 0 by definition. Symmetric probability density functions are associated, for instance, with quadratic potentials (Eq. C.10) that are typical of bifurcation normal forms under white noise, for which (Gardiner, 1985)

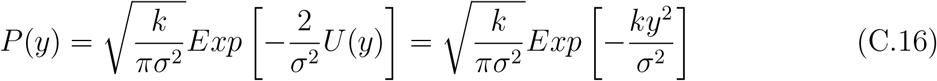

Consequently, the normal forms – in particular, the saddle-node – considered above are expected to display a flat skewness.

On the other hand, the integral C.16 may be non-zero, and even dependend on the drift parameter *k*, if *μ* ≠ 0 or if *P* (*y*) is asymmetrical. In the first case, solving Eq. C.16 yields (provided that Re[*k*] *>* 0):

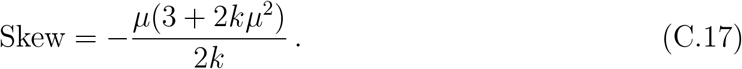

In this case, as *k* → 0, the skewness is expected to increase, potentially providing an early warning On the other hand, an asymmetric potential can be obtained in case of multiplicative noise (Gardiner, 1985; Sharma et al., 2016). Depending on the specific form, it may be possible to observe increasing trends associated to EWS, but they may be system-specific and not generalisable. In this sense, there is no ambiguity between the results of Guttal and Jayaprakash (2008) and Dai et al. (2012): they were studying systems with different properties, using an indicator that is not particularly performing and generalisable.

As for the kurtosisn, in case of *μ* = 0 (typical white noise), kurtosis = 3Var^2^. This can be obtained by solving Eq. C.16. If *μ* ≠ 0, or for other exotic noise forms, and if Re[*k*] *>* 0:

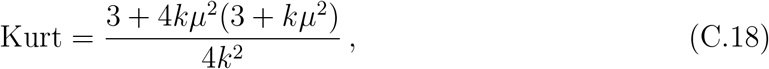

whose leading term for 0 *< k <* 1 still equals Var^2^. Hence, the variance is already representative of higher moments, which are not expected to improve EWS unless system-specific noise and drift forms are considered. Note that, for both Eq. C.17 and Eq. C.18, the constant noise level *σ* is normalised to 1 for ease of notation.

### STAR Method C.5. Computational simulations

In all computer simulations of Eq. 15, *K* = 0.1 to set bistability. The analysis concentrates on the upper stable branch of the bifurcation diagram (Fig. 4, right) to compare with white noise results. In this case, multiplicative noise corresponds to intrinsic regulatory mechanisms (Hasty et al., 2000; Norman et al., 2015) rather than stochasticity due to small numbers (Gillespie, 2000). Simulations are performed in Matlab (R2021b) using the Milstein method with a time step of 0.01 (arbitrary units). For quasi-steady state simulations, distributional data for each *c* from far to close the bifurcation point are computed upon stable values of system’s state, after a transient.

The Milstein method runs Monte Carlo chains over Itô-Taylor expanded stochastic differential equations for any variable *z*, up to second order:

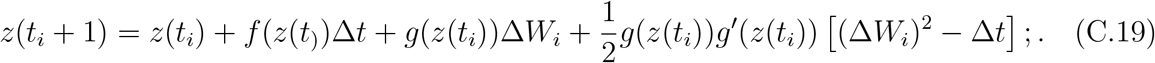

It better converges to the true Itô integral and was proven to have improved accuracy (Bayram et al., 2018). When *g*(*z*(*t*)) = *const* (only additive noise without state-dependency), it is equivalent to the common Euler-Maruyama scheme.

Setting simulation parameters of noise intensity and distance to critical points require understanding their reciprocal scales. To do so, we employ a methodology introduced in (Kuehn, 2011; Proverbio et al., 2022b), that is, to look for significant changes in the Kramers’ escape rates out of bistable potentials. The Kramers escape rate is (Gardiner, 1985)

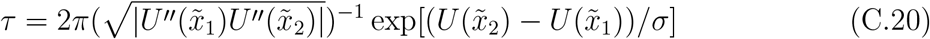

and measures the average expected rate of escape of multiple noisy particles from attracting wells. For any saddle-node bifurcation 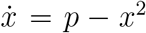 equipped with additive noise, 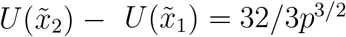 and 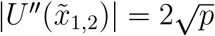. Hence,

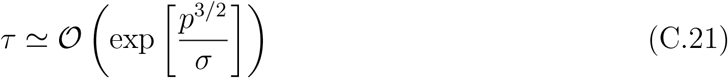

Comparable ranges of control parameters and noise levels are studied in (Proverbio et al., 2022b) and reproduced in Supplementary Figure S4. We use those results to distinguish two regimes, one where few noise-induced transitions might occur and another regime primarily determined by the approach to the bifurcation. We set values of *c* − *c*_0_ (distance from bifurcation point) and *σ* (noise intensity) accordingly, to span both regimes and see what changes when n-tipping becomes more frequent.

Finally, the statistical indicators are computed using their standard definitions, using their corresponding Matlab functions. For example, variance and Shannon entropy *H*_*S*_ are:

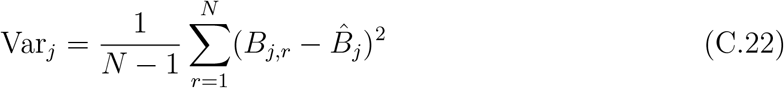

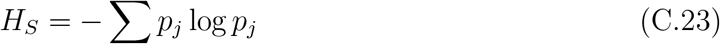

for any point *j* corresponding to a single parameter value, with *N* data *B* distributed around a mean value 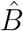 and probability density function *p*_*j*_. Other statistical moments and indicators can be computed similarly.

### STAR Method C.6. p-value assessment of significant increase and optimisation

By theory, an early warning signal is triggered when an increasing trend of suitable statistical indicators is observed (Scheffer et al., 2009). However, during real-time monitoring, it is often challenging to say whether a measured increase of mean values is significant or not, due to random fluctuations and uncertainties that may occur. If increasing trends are not quantified properly, spurious signals may be triggered (Boettiger and Hastings, 2012). For analysis performed using moving windows over time-series data, the Kendall’s *τ* score of monotonous increases have been proposed (Boettiger and Hastings, 2012; Proverbio et al., 2022a), as well as threshold of confidence intervals, with respect to baseline values (Drake and Griffen, 2010).

Since we work with distributional data, we propose to employ significance levels on Welch’s p-value scores (non-equal variances allowed between the populations), which relate to threshold in confidence intervals and are readily interpreteable (Proverbio et al., 2022b). They also allow to estimate the sensitivity to noise intensities and the expected lead parameter for detection or anticipation of critical transitions. The idea is to compare the full distributions at each parameter value *c* with a reference one, usually taken far from the bifurcation point and without n-tipping, and check whether they are significantly separated towards increasing values. The p-value scores are used to assess the significance. This method can still be sensitive to fluctuating scores (hence, a smoothing is employed), but it has the advantage of relying on a-prioristic values, e.g. significant p-value *p*_*sig*_ = 0.05. Of course, a p-value does not distinguish between increasing or decreasing trends: it is thus coupled with simple visualization of the direction of the trends.

Examples of the three methods are provided in Supplementary Figure S1.

Quantifying the significance of increasing trends is leveraged as follows: we extract at which value of the control parameter *c* the p-value crosses the significance threshold *p*_*sig*_ = 0.05 as a reference. Other common thresholds *p* = 0.1 or *p* = 0.01 can be used, yielding consistent results. When p-value *< p*_*sig*_, it means that an indicator has significantly increased more than the baseline, triggering a warning signal. Consider all *c*_*i*_ tested during the simulations, *i* = 1..*N* with *N* = (*c*_*max*_ − *c*_*min*_)*/*0.002; *c*_*max*_ and *c*_*min*_ are two arbitrary values greater and lower than the bifurcation value *c*_0_, within the bistable region, and 0.002 is the simulation step |*c*_*i*_ − *c*_*i*−1_|. Out of all *c*_*i*_, estimate *c*_*sig*_ = *c*_*j*_, where *j* is the first index at which p-value_*j*_ *< p*_*sig*_ stably, *i.e*., without considering small fluctuating values (for that, a smoothing is employed). This is performed for each indicator ℐ and each noise level *σ*.

Hence, the analysis estimates

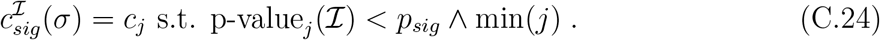

The optimisation problem described in the main text aims at maximising the combination of all 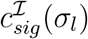 obtained at different noise levels *σ*_*l*_, so that the results are robust against a range of signal-to-noise ratios. As described in the main text, the analysis is complemented with a counter 𝒞to quantify how many tipping events occurred before the bifurcation point, for each *σ*.

A final comment regards the set of considered indicators ℐ. In principle, CV could be included among the as its performance improves in case of multiplicative noise (see Supplementary Figure S3. However, the optimisation procedure does not strongly select it, preferring the combinations in Fig. 5d. Hence, it has been removed altogether, to improve the computational speed when using more fine-grained steps for the grid search.

### STAR Method C.7. Data collection and analysis

Experimental data were collected and curated by the original study (Dai et al., 2012). We refer to it for details about the experimental protocols. The publicly available data correspond to ensemble of replicate populations, at each observation time corresponding to input dilution factors altering the environmental sucrose concentration. The eight dilution are 250, 500, 750, 1000, 1133, 1266, 1400 and 1600. Population densities were recorded by measuring optical density at 620 nm using a Thermo Scientific Multiskan FC microplate photometer. The values used in the analysis represent cell numbers, estimated from optical densities converted through calibration curves described in thee original publication. For each observation time, several statistical indicators were calculated over the ensembles as explained in the previous section.

The standard errors and confidence intervals of the indicators were given by bootstrap. In bootstrap, the replicates are resampled by combining the data over 5 days (observation lag for one dilution factor) into a single distribution. Resapling was performed by 50 to 1000 repetitions, to check the robustness of final p-values against bootstrapping hyperparameters and to confirm consistency with the original results. Since there are, on average, 60 data entries for each dilution factor value, we eventually employ bootstrapping with 50 repetitions, to avoid biases in the p-values due to random over-repetitions of some data.

The p-values to quantify significant increases in the distributions of indicators are calculated as described in STAR Method C.6, using the distribution at dilution factor 250 (the smallest and furthest from the bifurcation point) as baseline, and comparing all other distributions against it, making sure that the mean value was increasing before drawing conclusions.

## STAR Method D. Supplementary information titles and legends

**Figure S1: Quantitative definition of EWS**. Left: Example of looking for trends past thresholds of confidence intervals. In this case, past the 2*σ* interval (dashed line) over the uncertainty of the rightmost point, used as baseline far from the bifurcation value *c*_0_. Centre: Example of Kendall’s *τ* estimation. Compare the trends within two sliding windows. If the new one is monotonously increasing with respect to the old one, *τ >* 0, while no increase corresponds to *τ* = 0; the steeper the trend, the higher *τ*. Right: Example of p-value between two distributions corresponding to different parameter values: the baseline, corresponding to the rightmost *c*, and another generic *c*′. Each distribution corresponds to an average value of the statistical indicator (superimposed and shifted for visualization purposes). p-values’s significance can be checked with standard statistical methods, to assess whether the registered increase is significant or not. All figures use variance computed from simulations of Eq. 15 in main text, with *n* = 2, *K* = 0.1 and *σ* = 0.02. *c*_0_ is the critical value for bifurcation point.

**Figure S2: Trends of notable indicators before and after the bifurcation point** *c*_0_. It is displayed as a function of the control parameter *c* from Eq. 15 of main text. The increasing trends yield early warning signals. The violet ribbon represents confidence intervals of 2 standard deviations, estimated from repeated simulations. Indicators are: Variance, lag-1 Autocorrelation, Skewness, Kurtosis, Coefficient of Variation, Index of Dispersion, Shannon Entropy (*H*_*S*_). Note that some of them peak at the transition point, while others don’t due to noise-induced transitions altering their expected trends. All simulations are performed with white noise, *σ* = 0.012.

**Figure S3: Dependency of** *c*_*sig*_ **(Eq. C.24 of main text) for each considered indicator 𝒞 and noise intensity** *σ*. It is displayed with the corresponding counting 𝒞 of noise-induced transitions happening before the bifurcation point, at each noise intensity *σ*. Different multiplicative noise types are considered (*cf*. Eq. 17 of main text): a) *h*(*x*) = *x*. b) *h*(*x*) = *x*^2^. c) *h*(*x*) = *x*^2^*/*(1 + *x*^2^). Due to differing fluctuation types, the indicators have different performances in identifying the lead parameter. Conserved patterns are: entropy *H*_*S*_ is normally the best, particularly for high *σ*; Skew and Kurt perform poorly. AC(1) follows *H*_*S*_ closely, but with slightly lower *c*_*sig*_. Var and ID are normally worse that CV, as they are less sensitive to mean values. Notably, CV works better that in the case of white noise (compare with Fig. 5 of main text) but it still lags behind *H*_*S*_, particularly in case of low *σ*. Note that several n-tipping occur before the bifurcation point as *σ* increases, except for *h*(*x*) = *x*^2^*/*(1 + *x*^2^) that better buffers the system variability, as also noted in (Proverbio et al., 2022b). Particularly for this case, the main indicators provide anticipating signals (around *c*_*s*_*ig* ≥ 0.05) while n-tipping starts around *c* ≃ 0.02. In the other cases, the indicators are normally providing early warnings, except in the case *σ >* 0.046 for which they may just-on-time detect the few n-tipping events already happening.

**Figure S4: Kramers’ escape rate** *τ* **as a function of noise level** *σ* **and** *p* **(distance from bifurcation point)**. Its analytical form is in Eq. C.21 of main text of the main text. We use the boundary colored in yellow as a proxy to set commensurable magnitudes between control parameter and noise intensity in computer simulations.

## References

Aihara, K., Liu, R., Koizumi, K., Liu, X., Chen, L., 2022. Dynamical network biomarkers: Theory and applications. Gene 808, 145997. DOI https://doi.org/10.1016/j.gene.2021.145997

Allen, L. J. S., 2010. An introduction to stochastic processes with applications to biology. CRC press. DOI https://doi.org/10.1201/b12537

Alon, U., 2006. An introduction to systems biology: design principles of biological circuits. CRC press. DOI https://doi.org/10.1201/9781420011432

Andrecut, M., Halley, J. D., Winkler, D. A., Huang, S., 2011. A general model for binary cell fate decision gene circuits with degeneracy: Indeterminacy and switch behavior in the absence of cooperativity. PLoS ONE 6 (5), e19358. DOI https://doi.org/10.1371/journal.pone.0019358

Angeli, D., Ferrell, J. E., Sontag, E. D., 2004. Detection of multistability, bifurcations, and hysteresis in a large class of biological positive-feedback systems. Proceedings of the National Academy of Sciences 101 (7), 1822–1827. DOI https://doi.org/10.1016/j.sysconle.2003.08.003

Antoniou, D., Schwartz, S. D., 2011. Protein dynamics and enzymatic chemical barrier passage. The Journal of Physical Chemistry B 115 (51), 15147–15158. DOI https://doi.org/10.1021/jp207876k

Ashwin, P., Wieczorek, S., Vitolo, R., Cox, P., 2012. Tipping points in open systems: bifurcation, noise-induced and rate-dependent examples in the climate system. Philosophical Transactions of the Royal Society A 370 (1962), 1166–1184. DOI https://doi.org/10.1098/rsta.2011.0306

Ashwin, P., Zaikin, A., 2015. Pattern selection: The importance of “how you get there”. Biophysical Journal 108 (6), 1307–1308. DOI https://doi.org/10.1016/j.bpj.2015.01.036

Bayram, M., Partal, T., Buyukoz, G. O., 2018. Numerical methods for simulation of stochastic differential equations. Advances in Difference Equations 2018 (1), 1–10. DOI https://doi.org/10.1186/s13662-018-1466-5

Berglund, N., Gentz, B., 2006. Noise-induced phenomena in slow-fast dynamical systems: a sample-paths approach. Springer Science & Business Media. DOI https://doi.org/10.1007/1-84628-186-5

Boettiger, C., Hastings, A., 2012. Quantifying limits to detection of early warning for critical transitions. Journal of the Royal Society Interface 9 (75), 2527–2539. DOI https://doi.org/10.1098/rsif.2012.0125

Bonciolini, G., Ebi, D., Boujo, E., Noiray, N., 2018. Experiments and modelling of rate-dependent transition delay in a stochastic subcritical bifurcation. Royal Society Open Science 5 (3), 172078. DOI https://doi.org/10.1098/rsos.172078

Brett, T. S., Drake, J. M., Rohani, P., 2017. Anticipating the emergence of infectious diseases. Journal of The Royal Society Interface 14 (132), 20170115. DOI https://doi.org/10.1098/rsif.2017.0115

Bury, T. M., Sujith, R. I., Pavithran, I., Scheffer, M., Lenton, T. M., Anand, M., Bauch, C. T., 2021. Deep learning for early warning signals of tipping points. Proceedings of the National Academy of Sciences of the United States of America 118 (39), e2106140118. DOI https://doi.org/10.1073/pnas.2106140118

Carpenter, S. R., Cole, J. J., Pace, M. L., Batt, R., Brock, W. A., Cline, T., Coloso, J., Hodgson, J. R., Kitchell, J. F., Seekell, D. A., Smith, L., Weidel, B., 2011. Early warnings of regime shifts: A whole-ecosystem experiment. Science 332 (6033), 1079–1082. DOI https://doi.org/10.1126/science.1203672

Chen, L., Liu, R., Liu, Z. P., Li, M., Aihara, K., 2012. Detecting early-warning signals for sudden deterioration of complex diseases by dynamical network biomarkers. Scientific Reports 2, 18–20. DOI https://doi.org/10.1038/srep00342

Clements, C. F., Ozgul, A., 2018. Indicators of transitions in biological systems. Ecology Letters 21 (6), 905–919. DOI https://doi.org/10.1111/ele.12948

Cohen, A. A., Leung, D. L., Legault, V., Gravel, D., Blanchet, F. G., Côté, A.-M. C., Fülöp, T., Lee, J., Dufour, F., Liu, M., Nakazato, Y., 2022. Synchrony of biomarker variability indicates a critical transition: Application to mortality prediction in hemodialysis. iScience 25, 104385. DOI https://doi.org/10.1016/j.isci.2022.104385

Dai, L., Korolev, K. S., Gore, J., Carpenter, S. R., 2015. Relation between stability and resilience determines the performance of early warning signals under different environmental drivers. Proceedings of the National Academy of Sciences of the United States of America 112 (32), 10056–10061. DOI https://doi.org/10.1073/pnas.1418415112

Dai, L., Vorselen, D., Korolev, K. S., Gore, J., 2012. Generic indicators for loss of resilience before a tipping point leading to population collapse. Science 336 (6085), 1175–1177. DOI https://doi.org/10.1126/science.1219805

Dakos, V., Carpenter, S. R., van Nes, E. H., Scheffer, M., 2015. Resilience indicators: Prospects and limitations for early warnings of regime shifts. Philosophical Transactions of the Royal Society B 370 (1659), 1–10. DOI https://doi.org/10.1098/rstb.2013.0263

Del Vecchio, D., Dy, A. J., Qian, Y., 2016. Control theory meets synthetic biology. Journal of The Royal Society Interface 13 (120), 20160380. DOI https://doi.org/0.1098/rsif.2016.0380

Diks, C., Hommes, C., Wang, J., 2019. Critical slowing down as an early warning signal for financial crises? Empirical Economics 57 (4), 1201–1228. DOI https://doi.org/10.1007/s00181-018-1527-3

Dmitriev, A., Dmitriev, V., Sagaydak, O., Tsukanova, O., 2017. The Application of Stochastic Bifurcation Theory to the Early Detection of Economic Bubbles. Procedia Computer Science 122, 354–361. DOI https://doi.org/10.1016/j.procs.2017.11.380

Drake, J. M., Griffen, B. D., 2010. Early warning signals of extinction in deteriorating environments. Nature 467 (7314), 456–459. DOI https://doi.org/10.1038/nature09389

Drijfhout, S., Bathiany, S., Beaulieu, C., Brovkin, V., Claussen, M., Huntingford, C., Scheffer, M., Sgubin, G., Swingedouw, D., 2015. Catalogue of abrupt shifts in Intergovernmental Panel on Climate Change climate models. Proceedings of the National Academy of Sciences of the United States of America 112 (43), E5777–E5786. DOI https://doi.org/10.1073/pnas.1511451112

Dudney, J., Suding, K. N., 2020. The elusive search for tipping points. Nature Ecology & Evolution 4 (11), 1449–1450. DOI https://doi.org/10.1038/s41559-020-1273-8

Dunlop, M. J., Cox, R. S., Levine, J. H., Murray, R. M., Elowitz, M. B., 2008. Regulatory activity revealed by dynamic correlations in gene expression noise. Nature Genetics 40 (12), 1493–1498. DOI https://doi.org/10.1038/ng.281

Eugenio, M., Karp, R. L., Guo, G., Robson, P., Hart, A. H., Trippa, L., Yuan, G. C., 2014. Bifurcation analysis of single-cell gene expression data reveals epigenetic landscape. Proceedings of the National Academy of Sciences of the United States of America 111 (52), E5643–E5650. DOI https://doi.org/10.1073/pnas.1408993111

Feng, S., Sáez, M., Wiuf, C., Feliu, E., Soyer, O. S., 2016. Core signalling motif displaying multistability through multi-state enzymes. Journal of the Royal Society Interface 13 (123). DOI https://doi.org/10.1098/rsif.2016.0524

Ferrell Jr, J. E., Pomerening, J. R., Kim, S. Y., Trunnell, N. B., Xiong, W., Huang, C.-Y. F., Machleder, E. M., 2009. Simple, realistic models of complex biological processes: positive feedback and bistability in a cell fate switch and a cell cycle oscillator. FEBS letters 583 (24), 3999–4005. DOI https://doi.org/10.1016/j.febslet.2009.10.068

Gao, J., Barzel, B., Barabási, A. L., 2016. Universal resilience patterns in complex networks. Nature 530 (7590), 307–312. DOI https://doi.org/10.1038/nature16948

Gardiner, C. W., 1985. Handbook of stochastic methods - for Physics, Chemistry and the Natural Sciences. Springer Berlin. DOI https://doi.org/10.1007/978-3-662-02452-2

Ghaffarizadeh, A., Flann, N. S., Podgorski, G. J., 2014. Multistable switches and their role in cellular differentiation networks. BMC bioinformatics 15 (7), 1–13. DOI https://doi.org/10.1186/1471-2105-15-S7-S7

Gillespie, D. T., 2000. Chemical Langevin equation. Journal of Chemical Physics 113 (1), 297–306. DOI https://doi.org/10.1063/1.481811

Guttal, V., Jayaprakash, C., 2008. Changing skewness: An early warning signal of regime shifts in ecosystems. Ecology Letters 11 (5), 450–460. DOI https://doi.org/10.1111/j.1461-0248.2008.01160.x

Haragus, M., Iooss, G., 2010. Local Bifurcation, Center Manifolds and Normal Forms in Infinte-Dimensional Dynamical Systems. Springer Science & Business Media.

Hasty, J., Pradines, J., Dolnik, M., Collins, J. J., 2000. Noise-based switches and amplifiers forgene expression. Proceedings of the National Academy of Sciences of the United States of America 97 (5), 2075–2080. DOI https://doi.org/10.1073/pnas.040411297

Hirota, M., Holmgren, M., Van Nes, E. H., Scheffer, M., 2011. Global Resilience of Tropical Forest. Science 334 (October), 232–235. DOI https://doi.org/10.1126/science.1210657

Holling, C. S., 1996. Engineering resilience versus ecological resilience. Engineering within ecological constraints 31 (1996), 32. DOI https://doi.org/10.17226/4919

Horsthemke, W., Lefever, R., 1984. Noise-induced transitions in physics, chemistry, and biology. Noise-induced transitions: theory and applications in physics, chemistry, and biology, 164–200. DOI https://doi.org/10.1007/3-540-36852-37

Izhikevich, E. M., 2007. Dynamical systems in neuroscience: the geometry of excitability and bursting. MIT press. DOI https://doi.org/10.7551/mitpress/2526.001.0001

Khas’minskii, R. Z., 1966. A limit theorem for the solutions of differential equations with random right-hand sides. Theory of Probability & Its Applications 11 (3), 390–406. DOI https://doi.org/10.1137/1111038

Kitano, H., 2004. Biological robustness. Nature Reviews Genetics 5 (11), 826–837. DOI https://doi.org/10.1038/nrg1471

Korolev, K. S., Xavier, J., Gore, J., 2014. Turning ecology and evolution against cancer. Nature Reviews Cancer, 1–10. DOI https://doi.org/10.1038/nrc3712

Kuehn, C., 2011. A mathematical framework for critical transitions: Bifurcations, fast–slow systems and stochastic dynamics. Physica D: Nonlinear Phenomena 240 (12), 1020–1035. DOI https://doi.org/10.1016/j.physd.2011.02.012

Kuehn, C., Bick, C., 2021. A universal route to explosive phenomena. Science Advances 7 (16), 1–7. DOI https://doi.org/10.1126/sciadv.abe3824

Kuehn, C., Lux, K., Neamtu, A., 2022. Warning Signs for Non-Markovian Bifurcations: Color Blindness and Scaling Laws. Proceedings of the Royal Society A 478 (2259), 20210740. DOI https://doi.org/10.1098/rspa.2021.0740

Kuznetsov, Y. A., 2013. Elements of applied bifurcation theory. Vol. 112. Springer Science & Business Media. DOI https://doi.org/10.1007/b98848

Lade, S. J., Gross, T., 2012. Early warning signals for critical transitions: a generalized modeling approach. PLoS computational biology 8 (2), e1002360. DOI https://doi.org/10.1371/journal.pcbi.1002360

Lang, J., Nie, Q., Li, C., 2021. Landscape and kinetic path quantify critical transitions in epithelial-mesenchymal transition. Biophysical Journal 120 (20), 4484–4500. DOI https://doi.org/10.1016/j.bpj.2021.08.043

Lenton, T. M., Livina, V. N., Dakos, V., Van Nes, E. H., Scheffer, M., 2012. Early warning of climate tipping points from critical slowing down: comparing methods to improve robustness. Philosophical Transactions of the Royal Society A 370 (1962), 1185–1204. DOI https://doi.org/10.1098/rsta.2011.0304

Liu, X. M., Xie, H. Z., Liu, L. G., Li, Z. B., 2009. Effect of multiplicative and additive noise on genetic transcriptional regulatory mechanism. Physica A: Statistical Mechanics and its Applications 388 (4), 392–398. DOI https://doi.org/10.1016/j.physa.2008.10.030

MacArthur, B. D., Ma’ayan, A., Lemischka, I. R., 2009. Systems biology of stem cell fate and cellular reprogramming. Nature reviews Molecular cell biology 10 (10), 672–681. DOI https://doi.org/10.1038/nrm2766

Maini, P., Myerscough, M., Winter, K., Murray, J., 1991. Bifurcating spatially heterogeneous solutions in a chemotaxis model for biological pattern generation. Bulletin of mathematical biology 53 (5), 701–719. DOI https://doi.org/10.1007/BF02461550

Matsumori, T., Sakai, H., Aihara, K., 2019. Early-warning signals using dynamical network markers selected by covariance. Physical Review E 100 (5), 1–9. DOI https://doi.org/10.1103/PhysRevE.100.052303

Mazzocchi, F., 2012. Complexity and the reductionism–holism debate in systems biology. Wiley Interdisciplinary Reviews: Systems Biology and Medicine 4 (5), 413–427. DOI https://doi.org/10.1002/wsbm.1181

Meisel, C., Kuehn, C., 2012. Scaling effects and spatio-temporal multilevel dynamics in epileptic seizures. PLoS ONE 7 (2). DOI https://doi.org/10.1371/journal.pone.0030371

Moejes, F. W., Matuszyńska, A., Adhikari, K., Bassi, R., Cariti, F., Cogne, G., Dikaios, I., Falciatore, A., Finazzi, G., Flori, S., et al., 2017. A systems-wide understanding of photosynthetic acclimation in algae and higher plants. Journal of Experimental Botany 68 (11), 2667–2681. DOI https://doi.org/10.1093/jxb/erx137

Mojtahedi, M., Skupin, A., Zhou, J., Castaño, I. G., Leong-Quong, R. Y., Chang, H., Trachana, K., Giuliani, A., Huang, S., 2016. Cell Fate Decision as High-Dimensional Critical State Transition. PLoS Biology 14 (12), 1–28. DOI https://doi.org/10.1371/journal.pbio.2000640

Moris, N., Pina, C., Arias, A. M., 2016. Transition states and cell fate decisions in epigenetic landscapes. Nature Reviews Genetics 17 (11), 693–703. DOI https://doi.org/10.1038/nrg.2016.98

Namachchivaya, N. S., Leng, G., 1990. Equivalence of stochastic averaging and stochastic normal forms. Journal of Applied Mechanics 57 (4), 1011–1017. DOI https://doi.org/10.1115/1.2897619

Navid Moghadam, N., Nazarimehr, F., Jafari, S., Sprott, J. C., 2020. Studying the performance of critical slowing down indicators in a biological system with a period-doubling route to chaos. Physica A: Statistical Mechanics and its Applications 544, 123396. DOI https://doi.org/10.1016/j.physa.2019.123396

Norman, T. M., Lord, N. D., Paulsson, J., Losick, R., 2015. Stochastic switching of cell fate in microbes. Annual review of microbiology 69, 381–403. DOI https://doi.org/10.1146/annurev-micro-091213-112852

O’Regan, S. M., Burton, D. L., 2018. How stochasticity influences leading indicators of critical transitions. Bulletin of mathematical biology 80 (6), 1630–1654. DOI https://doi.org/10.1007/s11538-018-0429-z

Ozbudak, E. M., Thattai, M., Lim, H. N., Shraiman, B. I., Van Oudenaarden, A., 2004. Multistability in the lactose utilization network of Escherichia coli. Nature 427 (6976), 737–740. DOI https://doi.org/10.1038/nature02298

Pavithran, I., Sujith, R. I., 2021. Effect of rate of change of parameter on early warning signals for critical transitions. Chaos 31 (1). DOI https://doi.org/10.1063/5.0025533

Proverbio, D., Kemp, F., Magni, S., Gonçalves, J., 2022a. Performance of early warning signals for disease re-emergence: A case study on COVID-19 data. PLOS Computational Biology 18 (3), e1009958. DOI https://doi.org/10.1371/journal.pcbi.1009958

Proverbio, D., Montanari, A. N., Skupin, A., Gonçalves, J., 2022b. Buffering variability in cell regulation motifs close to criticality. Physical Review E 106, L032402. DOI https://doi.org/10.1103/PhysRevE.106.L032402

Quail, T., Shrier, A., Glass, L., 2015. Predicting the onset of period-doubling bifurcations in noisy cardiac systems. Proceedings of the National Academy of Sciences 112 (30), 9358–9363. DOI https://doi.org/10.1073/pnas.1424320112

Rosso, O. A., Larrondo, H. A., Martin, M. T., Plastino, A., Fuentes, M. A., 2007. Distinguishing noise from chaos. Physical Review Letters 99 (15), 1–4. DOI https://doi.org/10.1103/PhysRevLett.99.154102

Sarkar, S., Sinha, S. K., Levine, H., Jolly, M. K., Dutta, P. S., 2019. Anticipating critical transitions in epithelial-hybrid-mesenchymal cell-fate determination. Proceedings of the National Academy of Sciences of the United States of America 116 (52), 26343–26352. DOI https://doi.org/10.1073/pnas.1913773116

Scheffer, M., Bascompte, J., Brock, W. A., Brovkin, V., Carpenter, S. R., Dakos, V., Held, H., Van Nes, E. H., Rietkerk, M., Sugihara, G., 2009. Early-warning signals for critical transitions. Nature 461 (7260), 53–59. DOI https://doi.org/10.1038/nature08227

Sharma, Y., Dutta, P. S., Gupta, A. K., 2016. Anticipating regime shifts in gene expression: The case of an autoactivating positive feedback loop. Physical Review E 93 (3), 1–13. DOI https://doi.org/10.1103/PhysRevE.93.032404

Shi, J., Li, T., Chen, L., 2016. Towards a critical transition theory under different temporal scales and noise strengths. Physical Review E 93 (3), 1–13. DOI https://doi.org/10.1103/PhysRevE.93.032137

Sidney, R. C., Dunlop, M., Elowitz, M. B., 2010. A synthetic three-color reporter framework for monitoring genetic regulation and noise. Journal of Biological Engineering 4 (10), 1–12. DOI https://doi.org/10.1186/1754-1611-4-10

Sornette, D., 2006. Critical phenomena in natural sciences. Springer Science & Business Media. DOI https://doi.org/10.1017/CBO9781107415324.004

Stanoev, A., Schröter, C., Koseska, A., 2021. Robustness and timing of cellular differentiation through population-based symmetry breaking. Development 148 (3), dev197608. DOI https://doi.org/10.1242/dev.197608

Strogatz, S. H., 2015. Nonlinear dynamics and chaos. CRC press. DOI https://doi.org/10.1201/9780429492563

Stumpf, P. S., Smith, R. C., Lenz, M., Schuppert, A., Müller, F.-J., Babtie, A., Chan, T. E., Stumpf, M. P., Please, C. P., Howison, S. D., et al., 2017. Stem cell differentiation as a non-markov stochastic process. Cell Systems 5 (3), 268–282. DOI https://doi.org/10.1016/j.cels.2017.08.009

Su, Y., Bintz, M., Yang, Y., Robert, L., Ng, A. H. C., Liu, V., Ribas, A., Heath, J. R., Wei, W., 2019. Phenotypic heterogeneity and evolution of melanoma cells associated with targeted therapy resistance. PLoS computational biology 15 (6), e1007034. DOI https://doi.org/10.1371/journal.pcbi.1007034

Taylor, J. R., 1997. An Introduction to Error Analysis. University Science Books, Mill Valley, California. DOI https://doi.org/10.1063/1.882103

Thompson, J. M. T., Sieber, J., 2011. Predicting climate tipping as a noisy bifurcation: a review. International Journal of Bifurcation and Chaos 21 (02), 399–423. DOI https://doi.org/10.1142/S0218127411028519

Trefois, C., Antony, P. M., Goncalves, J., Skupin, A., Balling, R., 2015. Critical transitions in chronic disease: Transferring concepts from ecology to systems medicine. Current Opinion in Biotechnology 34, 48–55. DOI https://doi.org/10.1016/j.copbio.2014.11.020

Tsimring, L. S., 2014. Noise in biology. Reports on Progress in Physics 77.2 (2), 026601. DOI https://doi.org/10.1088/0034-4885/77/2/026601.Noise

Tu, C., D’Odorico, P., Suweis, S., 2021. Dimensionality reduction of complex dynamical systems. iScience 24 (1), 101912. DOI https://doi.org/10.1016/j.isci.2020.101912

Van Kampen, N. G., 1992. Stochastic processes in physics and chemistry. Vol. 1. Elsevier. DOI https://doi.org/10.1016/B978-0-444-52965-7.X5000-4

Wang, J., Zhang, K., Xu, L., Wang, E., 2011. Quantifying the Waddington landscape and biological paths for development and differentiation. Proceedings of the National Academy of Sciences 108 (20), 8257–8262. DOI https://doi.org/10.1073/pnas.1017017108

Wang, R., Dearing, J. A., Langdon, P. G., Zhang, E., Yang, X., Dakos, V., Scheffer, M., 2012a. Flickering gives early warning signals of a critical transition to a eutrophic lake state. Nature 492 (7429), 419–422. DOI https://doi.org/10.1038/nature11655

Wang, X., Li, L., Cheng, Y., Liu, Q., 2012b. Construction of gene regulatory networks with colored noise. Neural Computing and Applications 21 (8), 1883–1891. DOI https://doi.org/10.1007/s00521-011-0584-8

Weber, M., Buceta, J., 2013. Stochastic stabilization of phenotypic states: the genetic bistable switch as a case study. PloS one 8 (9), e73487. DOI https://doi.org/10.1371/journal.pone.0073487

Weinans, E., Quax, R., van Nes, E. H., de Leemput, I. A., 2021. Evaluating the performance of multivariate indicators of resilience loss. Scientific Reports 11 (1), 1–11. DOI https://doi.org/10.1038/s41598-021-87839-y

Wieczorek, S., Ashwin, P., Luke, C. M., Cox, P. M., 2011. Excitability in ramped systems: The compost-bomb instability. Proceedings of the Royal Society A 467 (2129), 1243–1269. DOI https://doi.org/10.1098/rspa.2010.0485

Wilkat, T., Rings, T., Lehnertz, K., 2019. No evidence for critical slowing down prior to human epileptic seizures. Chaos 29 (9), 2–7. DOI https://doi.org/10.1063/1.5122759

Yasemi, M., Jolicoeur, M., 2021. Modelling cell metabolism: a review on constraint-based steady-state and kinetic approaches. Processes 9 (2), 322. DOI https://doi.org/10.3390/pr9020322

Zhang, H., Chen, Y., Chen, Y., 2012. Noise Propagation in Gene Regulation Networks Involving Interlinked Positive and Negative Feedback Loops. PLoS ONE 7 (12), 1–8. DOI https://doi.org/10.1371/journal.pone.0051840

Zhou, J. X., Aliyu, D. S., Aurell, E., Huang, S., 2012. Quasi-potential landscape in complex multi-stable systems. Journal of the Royal Society Interface 9 (77), 3539–3553. DOI https://doi.org/10.1098/rsif.2012.0434

