## Supplementary figures for "Systematic analysis and optimization of early warning signals for critical transitions"

### SI. Supplementary Figures

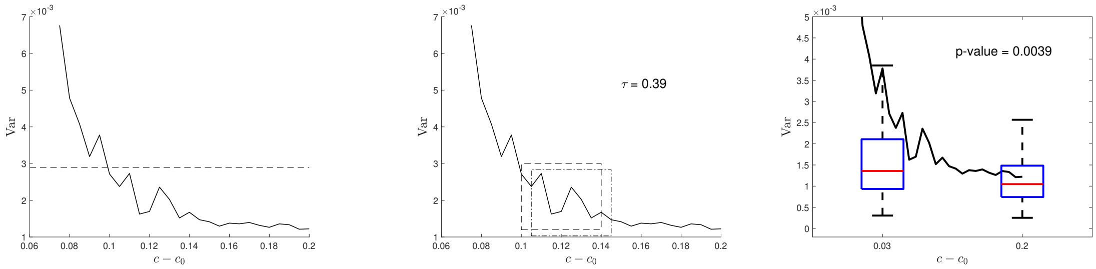


Figure S1: **Quantitative definition of EWS**. Left: Example of looking for trends past thresholds of confidence intervals. In this case, past the $2\sigma$ interval (dashed line) over the uncertainty of the rightmost point, used as baseline far from the bifurcation value $c_0$. Centre: Example of Kendall's $\tau$ estimation. Compare the trends within two sliding windows. If the new one is monotonously increasing with respect to the old one, $\tau > 0$, while no increase corresponds to $\tau = 0$; the steeper the trend, the higher $\tau$. Right: Example of p-value between two distributions corresponding to different parameter values: the baseline, corresponding to the rightmost $c$, and another generic $c'$. Each distribution corresponds to an average value of the statistical indicator (superimposed and shifted for visualization purposes). p-values's significance can be checked with standard statistical methods, to assess whether the registered increase is significant or not. All figures use variance computed from simulations of Eq. 15 in main text, with $n = 2$, $K = 0.1$ and $\sigma = 0.02$. $c_0$ is the critical value for bifurcation point.

*Correspondence:
`` (Daniele Proverbio)

### References

Proverbio, D., Montanari, A. N., Skupin, A., Gonçalves, J., 2022. Buffering variability in cell regulation motifs close to criticality. Physical Review E 106, L032402.
URL https://doi.org/10.1103/PhysRevE.106.L032402

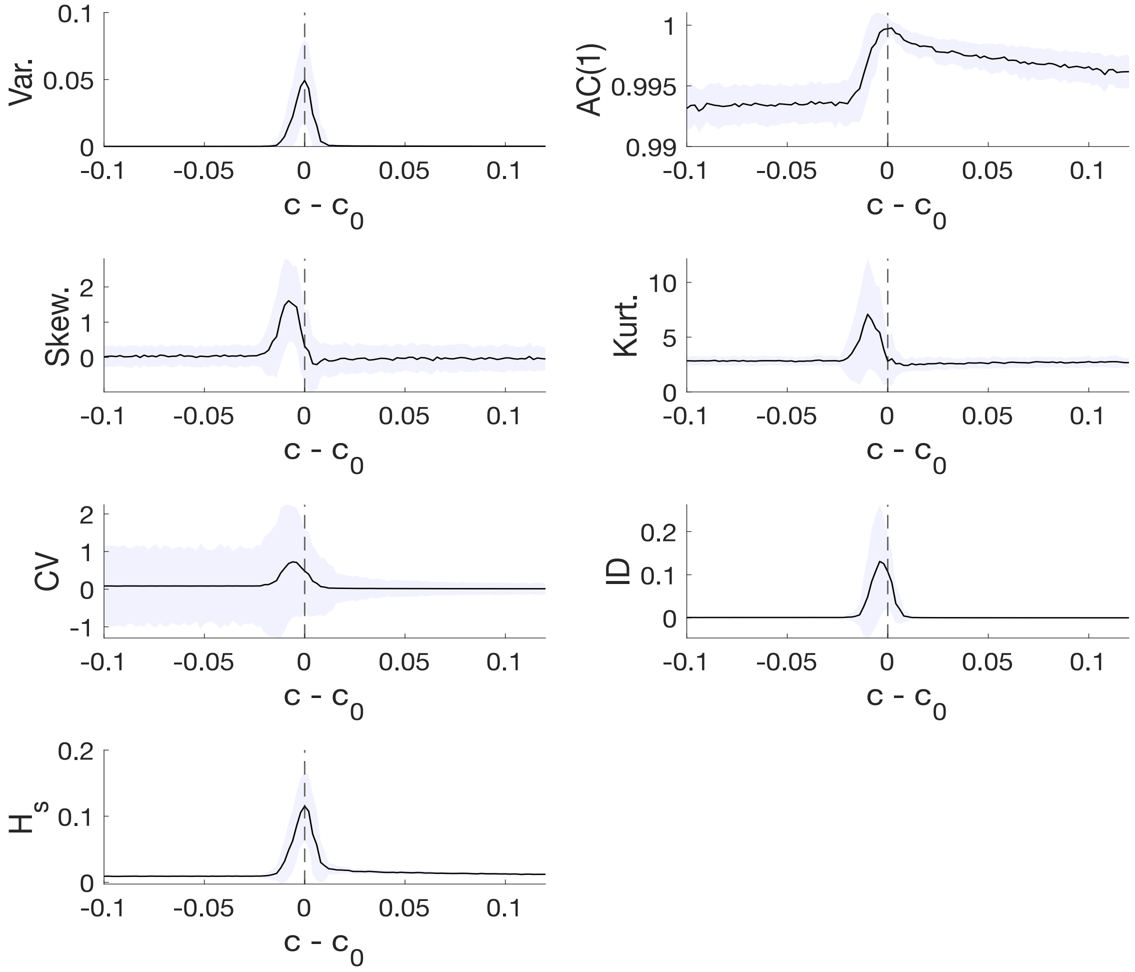


Figure S2: **Trends of notable indicators before and after the bifurcation point** $c_0$. It is displayed as a function of the control parameter $c$ from Eq. 15 of main text. The increasing trends yield early warning signals. The violet ribbon represents confidence intervals of 2 standard deviations, estimated from repeated simulations. Indicators are: Variance, lag-1 Autocorrelation, Skewness, Kurtosis, Coefficient of Variation, Index of Dispersion, Shannon Entropy ($H_S$). Note that some of them peak at the transition point, while others don't due to noise-induced transitions altering their expected trends. All simulations are performed with white noise, $\sigma = 0.012$.

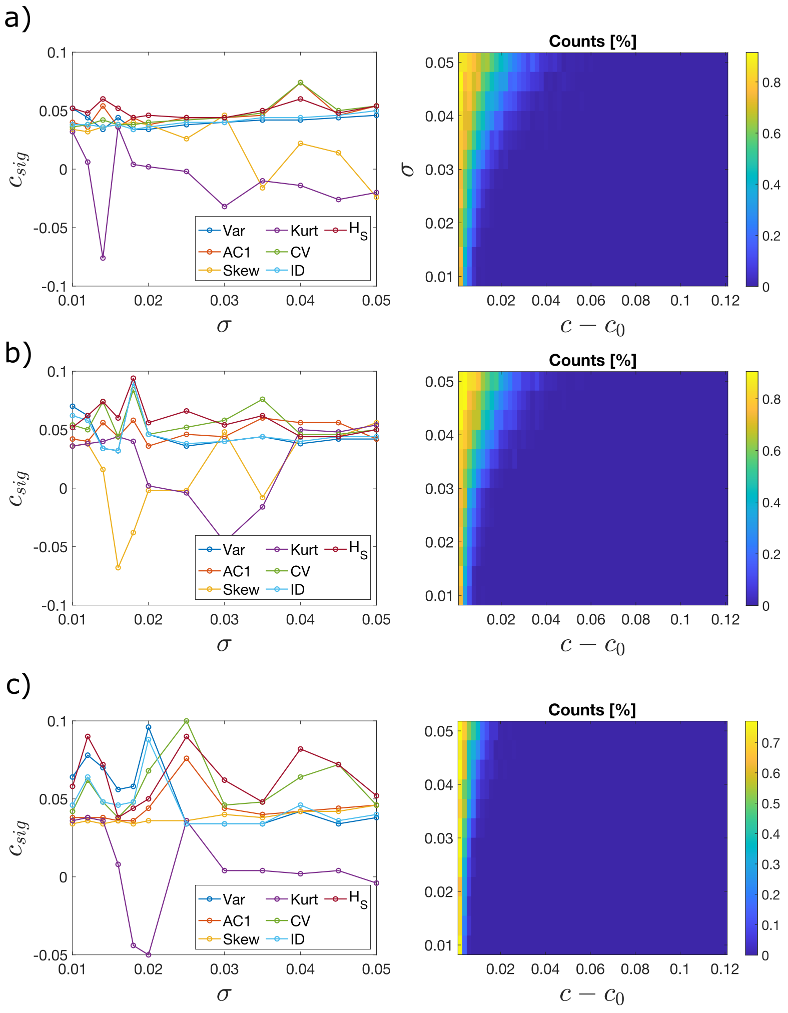


Figure S3: **Dependency of $c_{sig}$ (Eq. C.24 of main text) for each considered indicator $\mathcal{I}$ and noise intensity $\sigma$.** It is displayed with the corresponding counting $\mathcal{C}$ of noise-induced transitions happening before the bifurcation point, at each noise intensity $\sigma$. Different multiplicative noise types are considered (*cf.* Eq. 17 of main text): a) $h(x) = x$. b) $h(x) = x^2$. c) $h(x) = x^2/(1+x^2)$. Due to differing fluctuation types, the indicators have different performances in identifying the lead parameter. Conserved patterns are: entropy $H_S$ is normally the best, particularly for high $\sigma$; Skew and Kurt perform poorly. AC(1) follows $H_S$ closely, but with slightly lower $c_{sig}$. Var and ID are normally worse that CV, as they are less sensitive to mean values. Notably, CV works better that in the case of white noise (compare with Fig. 5 of main text) but it still lags behind $H_S$, particularly in case of low $\sigma$. Note that several n-tipping occur before the bifurcation point as $\sigma$ increases, except for $h(x) = x^2/(1+x^2)$ that better buffers the system variability, as also noted in (Proverbio et al., 2022). Particularly for this case, the main indicators provide anticipating signals (around $c_sig \geq 0.05$) while n-tipping starts around $c \simeq 0.02$. In the other cases, the indicators are normally providing early warnings, except in the case $\sigma > 0.046$ for which they may just-on-time detect the few n-tipping events already happening.

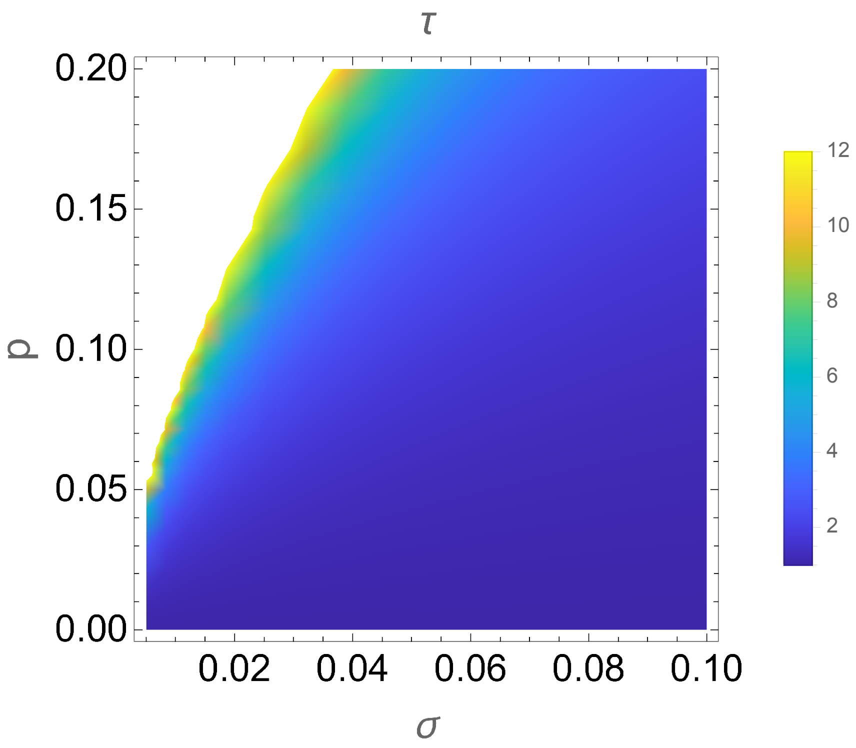


Figure S4: **Kramers' escape rate $\tau$ as a function of noise level $\sigma$ and $p$ (distance from bifurcation point)**. Its analytical form is in Eq. C.21 of main text of the main text. We use the boundary colored in yellow as a proxy to set commensurable magnitudes between control parameter and noise intensity in computer simulations.
